# A comparative analysis of cannabis and tobacco smoke exposure on human airway epithelial cell gene expression, immune phenotype, and response to formoterol and budesonide treatment

**DOI:** 10.1101/516294

**Authors:** Jennifer A. Aguiar, Ryan D. Huff, Wayne Tse, Martin R. Stampfli, Brendan J. McConkey, Andrew C. Doxey, Jeremy A. Hirota

## Abstract

Global recreational cannabis use is a potentially important public health issue that would benefit from experimental evidence to inform policy, regulations, and individual user practices. Comparative analyses between cannabis and tobacco smoke, the latter long reported to have negative impacts on respiratory health, may help provide context and provide clinically relevant evidence.

To address this unmet need we performed a comparative study between cannabis and tobacco smoke exposure in the Calu-3 human airway epithelial cells using concentration-response and pharmacological intervention study designs with outcome measurements of cell viability, epithelial cell barrier function, cytokine profile, and transcriptomics.

Our results demonstrate that cannabis smoke exposure reduces epithelial cell barrier function without impacting cell viability, accompanied by a cytokine profile associated with inflammation (elevated IL-6 and IL-8), barrier repair (elevated TGF-α and PDGF-AA) and suppressed antiviral immunity (decreased IP-10 and RANTES). Transcriptomic analyses revealed a cannabis smoke induced signature associated with suppressed antiviral genes and induction of oncogenic and oxidative stress pathways. Similar trends were observed for tobacco smoke exposure. A formoterol/budesonide intervention was unable to prevent cannabis smoke-induced reductions in antiviral pathways or normalize induction of oncogenic and oxidative stress responses.

Our results show striking similarities between cannabis and tobacco smoke exposure on impairing barrier function, suppressing antiviral pathways, potentiating of pro-inflammatory mediators, and inducing oncogenic and oxidative stress gene expression signatures. Furthermore, we demonstrate that an intervention with formoterol and budesonide is unable to completely normalized cannabis-induced responses. Collectively our data suggest that cannabis smoke exposure is not innocuous and may possess many of the deleterious properties of tobacco smoke, warranting additional studies to support public policy, government regulations, and individual user practices.

## INTRODUCTION

The United Nations World Drug Report estimates that over 180 million individuals use cannabis worldwide(1). With the decriminalization and legalization of cannabis in several states in the United States of America and across Canada in 2018, greater access to medical or recreational cannabis may lead to increased use of cannabis products. In Canada, the majority of cannabis is consumed by combustion with 94 and 89% of participants reporting this method of delivery in the 2017 and 2018 Canadian Cannabis Surveys, respectively(2, 3). The negative effects of *tobacco* smoke exposure on the lung and its airway epithelium are universally accepted (4–19). In comparison, the consequences of cannabis smoke exposure on lung health are less clear (20–29) and must receive additional attention to effectively inform public health policy, government regulations, and individual user practices.

Inhalation of smoke from combusted cannabis exposes the lungs to pharmacologically active ingredients, such as tetrahydrocannabinol (THC) and cannabidiol (CBD), as well as combustion products such as polycyclic aromatic hydrocarbons that are shared with biomass exposures, including tobacco(30). The psychoactive and immunomodulatory effects of cannabis have historically been attributed to THC and CBD, respectively, although an increasing body of evidence suggests complex interactions between the two (31–34). Habitual cannabis smoke exposure is associated with higher incidence of coughing, wheezing, chest tightness, and shortness of breath relative to non-smokers (20, 25–28); symptoms that are shared with tobacco smoking despite the difference in chemical composition. Comparative studies between cannabis and tobacco smoke are likely to reveal commonalities that are important in understanding the negative impacts that these exposures pose on lung health.

The lungs are in constant contact with harmful environmental agents, such as viruses and bacteria; yet we rarely show signs of infection(35, 36). Minimized infection is the result of a coordinated innate immune system in the lungs that begins with the mechanical barrier and immunological functions of the epithelium. Airway epithelial cells play a dominant role in creating a physical barrier between the external and internal environments in addition to producing mucus and antimicrobial peptides to trap and kill inhaled pathogens(36). If pathogens are capable of penetrating and overwhelming the defences provided by mucus and airway surface lining fluid, innate immune receptors on airway epithelial cells are poised to recognize molecular structures and trigger production and release of immune mediators. Despite this multi-tiered defence strategy, innate immune protection rendered by the airway epithelium can be compromised by tobacco smoke (4–6, 8–11) leading to increased susceptibility to bacterial or viral infection and potential for host pathology. Whether cannabis smoke exposure similarly impacts airway epithelial cell function and immune profile relevant in pathogen defence remains to be determined.

Tobacco research benefits from the availability of standardized research grade product that has been extensively characterized for composition and used in *in vitro* and *in vivo* exposure models (8, 10, 37–40). In contrast, cannabis research is currently limited by the lack of a standardized product, which may lead to an incomplete understanding of the consequences resulting from inhaled combusted smoke. The importance of using a cannabis strain with known chemical composition is emphasized by the proposed dependence of the immunomodulatory effects of cannabis on CBD concentrations (31–34, 41–43). Whether variation in THC and CBD composition between cannabis strains may differentially impact immune responses in the lung, including those orchestrated by airway epithelial cells, remains to be explored. A step forward in performing cannabis exposure related science should include reporting of the chemical composition of THC and CBD where present.

To manage chronic lung diseases an individual may be prescribed a long acting beta-agonist/glucocorticoid (LABA/GCS) combination treatment (44–47). LABA/GCS are transcriptionally active in airway epithelial cells resulting in broad anti-inflammatory activities that are believed to be important in controlling infections and chronic lung disease daily management and to prevent exacerbations (48–50). Importantly, the efficacy of LABA/GCS therapies may be compromised by tobacco smoke exposure (8, 13). It remains to be determined whether cannabis smoke exposure similarly compromises the efficacy of LABA/GCS anti-inflammatory activities. As cannabis use becomes more universally accepted, it is important to understand the effects that cannabis use has on airway health and the interventions used to manage lung inflammation and disease.

Based on the existing knowledge, we performed a comparative study between cannabis and tobacco smoke exposure in the Calu-3 human airway epithelial cell line. We performed a concentration-response study (7 concentrations) with cannabis and tobacco smoke to define airway epithelial cell viability, barrier function, and cytokine profile. Using a 10% smoke extract dilution informed by our concentration-response study, we performed a LABA/GCS intervention study with formoterol and budesonide and performed outcome measurements of airway epithelial cell viability, barrier function, cytokine profile, and RNA-sequencing based transcriptomics. We hypothesized that exposure of airway epithelial cells to cannabis smoke would negatively impact function and the profile of immune mediators, which are important features of the epithelium that protect the lung from inhaled pathogens. Our results demonstrate that cannabis smoke exposure reduces epithelial cell barrier function without impacting cell viability, accompanied by a cytokine profile associated with inflammation (elevated IL-6 and IL-8), barrier repair (elevated TGF-α and PDGF-BB) and suppressed antiviral immunity (decreased IP-10 and RANTES). Our LABA/GCS intervention reveals that this class of anti-inflammatories is able to suppress inflammation (IL-6 and IL-8) and improve barrier function, while further attenuating antiviral responses. In addition, transcriptomic signatures associated with oxidative stress were unaffected by LABA/GCS. Strikingly, broadly speaking, both cannabis and tobacco smoke exposure induced the same responses in outcome measurements assessed, suggesting that the negative health impacts attributed to tobacco may be similarly induced by cannabis.

## MATERIALS AND METHODS

### Preparation of tobacco cigarette and marijuana smoke extracts

Cannabis smoke extract (CSE) and tobacco smoke extract (TSE) conditioned media were freshly prepared via minor modifications to previously published methods (8–10, 13, 38). For generation of the TSE, a Kentucky Research Grade Cigarette (Lot: 3R4F) (~0.7g of dried tobacco leaves) was used. For generation of the CSE, cannabis from Dr. Jonathan Page (University of British Columbia, Vancouver, British Columbia, Canada) (13% THCA strain (w/w), with 0.18% THC, 0.35% THCVA, and 0.18% CBGA; ~0.7g dried cannabis rolled with cardboard filters) was used. To prepare the smoke-conditioned media, either 1 cannabis cigarette or 1 tobacco cigarette was smoked into 4 ml of HEPES buffered Eagle’s Minimal Essential Medium (EMEM). Crude smoke extracts were filtered using a 0.22μm filter. Extracts were standardized by measuring absorbance and diluting with fresh medium to reach a desired dilution (OD260nm = 0.4045*dilution factor, 10% dilution = 0.04045 OD260nm). A single batch of CSE and TSE was generated, aliquoted, and stored at −80°C and used for all subsequent experiments.

### Epithelial cell culture and drugs

Calu-3 cells, an immortalized human lung adenocarcinoma cell line, were obtained from ATCC (HTB-55, Lot: 61449062) and maintained in Eagle’s Minimum Essential Medium (EMEM, Millipore Sigma) supplemented with 10mM HEPES (Millipore Sigma), 10% FBS (Millipore Sigma), and Antibiotic-Antimycotic (Gibco®, ThermoFisher Scientific). Calu-3 cells were cultured in vented 75cm^2^ tissue culture flasks (Sarstedt) and used between passages 10-20. For exposure experiments 1×10^6^ or 2×10^5^ Calu-3 cells were seeded onto either 4.7cm^2^ or 0.3cm^2^ polyester Transwell^®^ permeable supports with a pore size of 0.4μm (Corning) and grown for 20 days to allow the Calu-3 cells to semi-differentiate into a pseudo-stratified epithelium as shown in previous studies(51). FBS was removed from Calu-3 cultures 24 hours prior to the start of the exposures. For exposures, fresh FBS free EMEM culture media was added to basal chambers and CSE or TSE media diluted with fresh FBS free EMEM culture media was added to the apical chambers for 24 hours. For intervention conditions, Budesonide 100nM (Cayman Chemical) and Formoterol 10nM (Cayman Chemical) was added to the exposure medias before being applied to Calu-3 cells for 24 hours.

### Cell viability assay

Cell viability was assessed using a Pierce™ LDH Cytotoxicity Assay Kit (ThermoFisher Scientific) according to the manufacturer’s instructions.

### Barrier function assessment (TEER)

Transepithelial Electrical Resistance (TEER) was measured using a Millicell ERS-2 Voltohmmeter with a STX01 electrode (Millipore Sigma) on Calu-3 cells grown on a polyester Transwell^®^ permeable support insert with a growth area of either 0.3cm^2^ (pharmacological intervention studies) or 4.7cm^2^ (RNA sequencing studies) with a pore size of 0.4μm (Corning). Resistance (Ohms) across the monolayer submerged in FBS free EMEM culture media was measured just prior to exposure as well as 24 hours post-exposure and multiplied by the growth area of the inserts (Ohms*cm^2^).

### Cytokine Assays

Following 24 hours of exposure, cell supernatants were collected, centrifuged, aliquoted, and shipped to Eve Technologies (Calgary, Alberta, Canada) to be analyzed with either a 42-plex or 65-plex human cytokine/chemokine panel. Concentrations were measured in pg/mL. Values that were below the limit of detection ½(LOD) were floored so as to equal ½(LOD). Two mediators from the LABA/GCS intervention study (ENA-78 and VEGF-A) had values exceeding the LOD and were removed from the analysis to avoid issues with signal saturation.

### RNA sequencing analysis

Total RNA was extracted using a RNeasy Plus Kit (QIAGEN, Valencia, CA) according to the manufacturer’s instructions. cDNA was prepared at The Centre for Applied Genomics at the Hospital for Sick Children (Toronto, Ontario, Canada). Samples were sequenced on the Illumina HiSeq 2500 instrument with 125 bp paired-end reads to a minimum depth of 30 million reads per sample. Reads were de-multiplexed and trimmed at The Centre for Applied Genomics and BCL files generated from the Illumina sequencer were converted to fastq files prior to our receivement of the reads.

After quality control using FastQC (v.0.11.7) and Prinseq (v.0.20.4), sequences were aligned to the human reference genome (hg19) using HISAT2 (v.2.1.0) and assembled into full transcriptomes using StringTie (v.1.3.3b). Samtools (v.1.9) was used to convert and sort HISAT2 output into sorted bam files for use by StringTie. StringTie was also used to calculate transcript abundances for downstream differential expression analysis using the Ballgown package in R (v. 3.4.3.) which provides p values, FDR-adjusted *p* values (*q* values), and fold change values for all genes in each comparison. A snakemake-based pipeline called hppRNA (v.1.3.3) was used to combine the above steps into a more simplistic work-flow.

### Functional Enrichment and Pathway Analyses

Lists of significantly up-regulated and down-regulated genes shared between CSE and TSE (as determined by Ballgown, FDR *q*-value < 0.05) were submitted to EnrichR to identify significantly enriched pathways and functional ontologies. Terms were ranked within ontologies by combined score which EnrichR calculates by taking the log of the *p* value derived from the Fisher exact test and multiplying that by the z-score of the deviation of the expected rank. Expected rank (adjusted *p* value) was calculated by EnrichR by running the Fisher exact test for many random gene sets in order to compute a mean rank and standard deviation from the expected rank for each term in the gene set library.

### Effects of Smoke Exposure and Formoterol/Budesonide Intervention on Oxidative Stress, Benz-a-pyrene-related, Pro-inflammatory, and Anti-viral Genes

Genes of interest with respect to oxidative stress, benzo-a-pyrene-induced pathways, inflammation, and anti-viral capability were selected from literature and used to assess the effect cannabis smoke exposure may have on these pathways compared to tobacco and whether formoterol/budesonide intervention successfully attenuates observed changes. Log_2_(mean FPKM) for the most abundant transcript of each gene was compared across growth environments to identify significant differential expression.

### Fluorescence Microscopy and Oxidative Stress Analysis

Cell-permeable redox-sensitive fluorescent dyes were used to assess oxidative stress in Calu-3 cells seeded into 96-well clear bottom black walled plates (Corning) at 8.0×10^4^cells/well. H_2_O_2_ (ThermoFisher Scientific) at a concentration of 0.5mM in FBS free EMEM culture media was used as a positive control. For H_2_DCFDA (ThermoFisher Scientific) assays, Calu-3 cells were washed with HBSS with Ca^2+^ Mg^2+^ (ThermoFisher Scientific) prior to treatment with 15μM H_2_DCFDA diluted in HBSS with Ca^2+^ Mg^2+^ for 30 minutes. Treated Calu-3 cells were then washed three times with HBSS with Ca^2+^ Mg^2+^ and exposed to smoke extracts as previously described for 24 hours. For CellROX Green Reagent assays (ThermoFisher Scientific), Calu-3 cells were exposed to smoke extracts as previously described for 24 hours, washed with HBSS with Ca^2+^ Mg^2+^ three times, incubated with 5μM CellROX Green Reagent and 1μg/mL Hoechst 3342 (BD Biosciences) for 30 minutes, and then washed three times with HBSS with Ca^2+^ Mg^2+^. Relative fluorescent intensity of H_2_DCFDA (488nm excitation/525nm emission), CellROX (485nm excitation/520nm emission) and Hoechst (360nm excitation/470nm emission) was measured using a SpectraMaxi3x (Molecular Devices) fluorescent plate reader. Calu-3 cells treated with CellROX and Hoechst were imaged within 30 minutes using an EVOS FL Auto Cell Imaging System.

### Statistical Analysis for *In Vitro* Experiments

Significant changes in cell viability, TEER, and cytokines were identified through permutation ANOVA followed by Tukey Honest Significant Difference (HSD) post-hoc test using the “lmPerm” package in R (v. 3.4.3.). For all analyses, differences were considered statistically significant when adjusted *p* values are less than 0.05. For all experiments n=4 independent experimental trials.

## RESULTS

### Concentration-Response analysis of cannabis smoke exposure on airway epithelial cell viability, barrier function, and immune profile

The impact of cannabis smoke exposure on airway epithelial cell viability, barrier function, and immune profile relative to tobacco smoke were examined using a concentration-response experiment design using dilutions of smoke-conditioned cell culture media (**Figure 1**).

**Figure 1.**
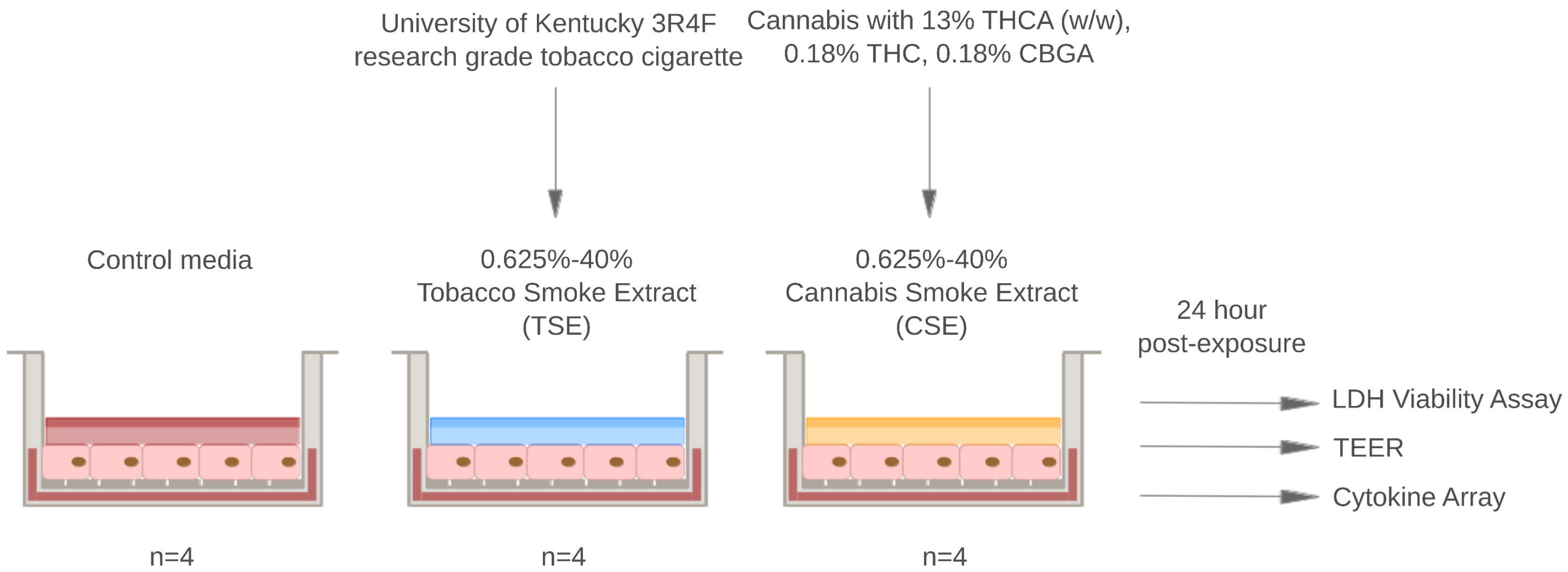
Concentration-Response Experiment Work-Flow. The impact of cannabis smoke exposure on airway epithelial cell barrier viability, barrier function, and immune profile relative to tobacco smoke were examined using a concentration-response experiment design using dilutions of smoke-conditioned cell culture media. Dilutions ranged from 0.625%-40% with 0% (untreated) cells as the control. Calu-3 cells were exposed in replicates of 4 per dilution/smoke extract for 24 hours before downstream analysis via LDH viability assay, TEER, and cytokine array.

### Cannabis smoke exposure does not impact airway epithelial cell viability

Cell viability as measured by lactate dehydrogenase (LDH) levels in cell culture supernatant were quantified and revealed no change in cell viability for cannabis smoke exposure at any concentration examined (0.625-40% diluted conditioned media) (**Figure 2A**). Similarly, tobacco smoke exposure had minimal impact on cell viability with the only increase in LDH measured at 40% dilution. Our results suggest that cannabis smoke extract has minimal impact on cell viability in our model.

**Figure 2.**
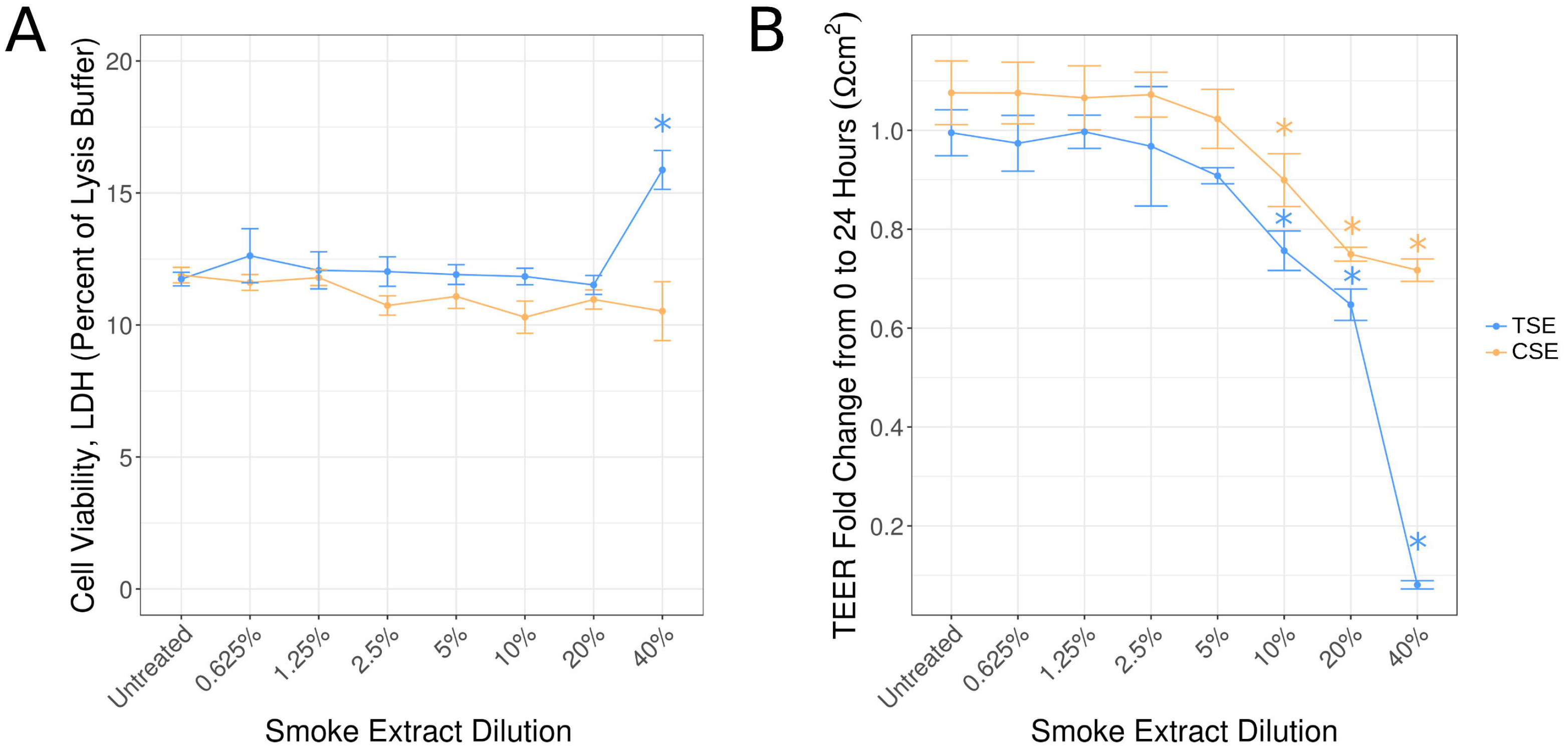
Effect of cannabis smoke exposure on airway epithelial cell viability and barrier function. Calu-3 cells were exposed to 7 concentrations of CSE or TSE for 24 hours. ***A*)** Cell viability was assessed via LDH. ***B*)** Fold change TEER measured at 24 hours post-exposure compared to untreated cells. CSE (orange), TSE (blue) * = p<0.05 relative to control untreated - Tukey HSD.

### Cannabis smoke exposure reduces airway epithelial cell barrier function in a concentration-dependent manner

Epithelial cell barrier function was measured by analyzing trans-epithelial electrical resistance (TEER) in transwell cultures reveling a concentration-dependent reduction for cannabis smoke exposures with significant decreases observed at 10, 20 and 40% dilutions (*p*<0.05 **– Figure 2B**). Similarly, tobacco smoke exposure induced a concentration-dependent reduction in epithelial cell barrier function above 10% dilutions with the greatest decrease in all experiments observed when cell viability was compromised (40%).

### Cannabis smoke exposure shifts airway epithelial cell immune profile towards repair, inflammation, and suppression of antiviral immunity

The impact of 40% tobacco smoke extract on cell viability with a major impact on epithelial cell barrier function suggested that this concentration was toxic and should be avoided for future studies. For this reason, all subsequent concentration-response analyses of immune profiles used the 0.625%-20% dilution range.

The impact of cannabis smoke exposure on immune profile was determined using a human multiplex cytokine array of 42-mediators. Our *a priori* hypothesis was that cannabis smoke extract would increase production of mediators important in epithelial cell barrier repair (TGF-α and PDGF-AA), decrease anti-viral mediators (IP-10 and RANTES), and increase pro-inflammatory mediators (IL-8 and IL-6). Data from additional cytokines analyzed are available in the Online Supplement (**Supplemental Table 1**).

In the context of epithelial cell barrier repair mediators, cannabis smoke induced a concentration-dependent increase in TGF-α and PDGF-AA that was significant at 20% dilution (*p*<0.05 – **Figure 3A-B**). Tobacco smoke exposure similarly increased TGF-α and PDGF-AA that was significant at 10% dilution.

**Figure 3.**
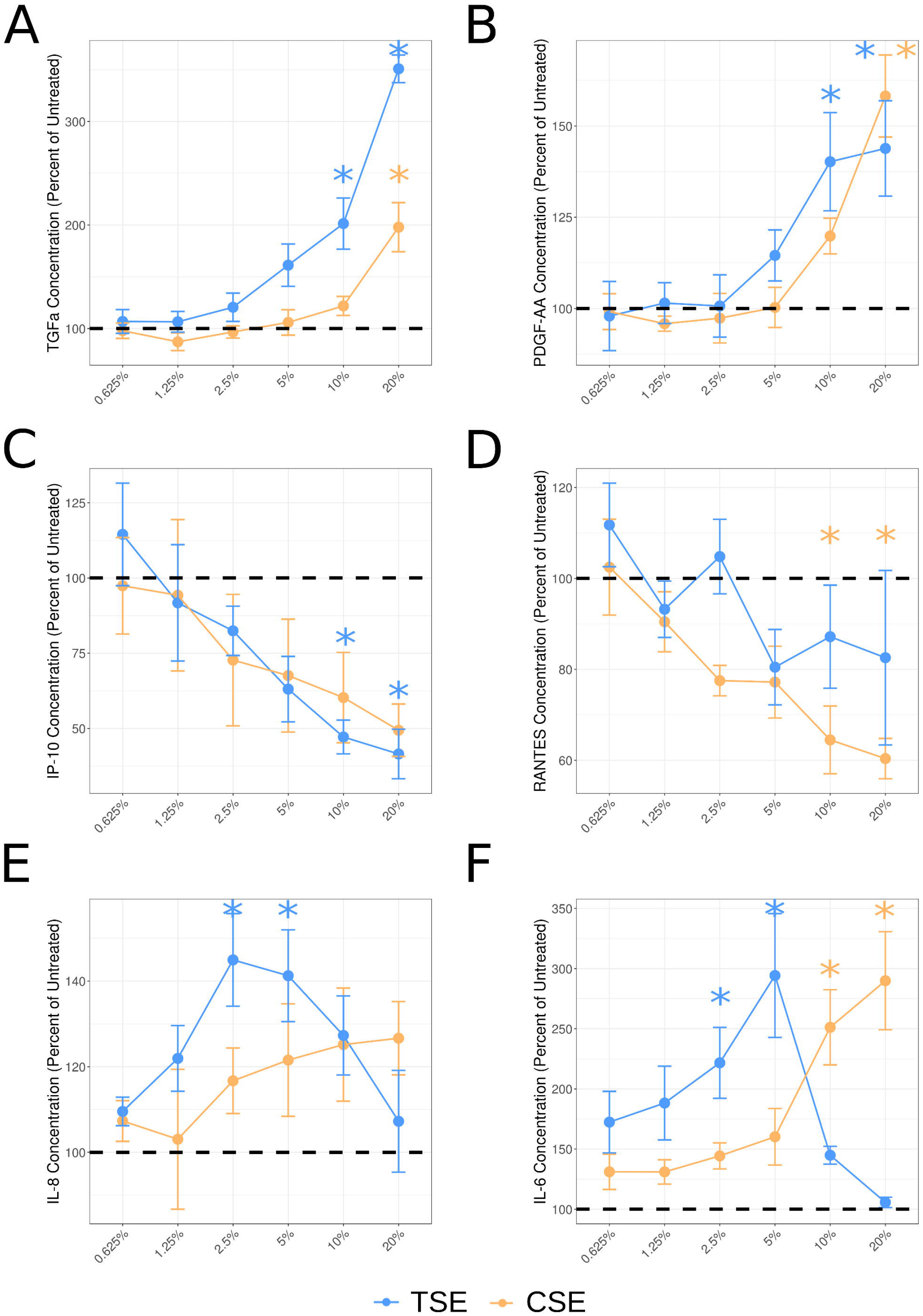
Effect of cannabis smoke exposure on select cytokine production. Dilutions of CSE (orange) and TSE (blue) were compared to untreated Calu-3 cells with respect to cytokine production (pg/mL). ***A*)** TGF-α ***B*)** PDGF-AA, ***C*)** IP-10 ***D*)** RANTES, ***E*)** IL-8, and ***F*)**IL-6. * = p<0.05 relative to control untreated - Tukey HSD.

In the context of antiviral immune mediators, we observed a concentration-dependent decrease in RANTES in response to cannabis smoke exposure that was significant beyond 10% (*p*<0.05 – **Figure 3D)**while a trend was observed for IP-10 (p>0.05 – **Figure 3C**). Tobacco smoke exposure induced a similar trend for both mediators with a significant decrease in IP-10 beyond 10% (*p*<0.05 – **Figure 3C**) and a trend for a reduction in RANTES/CXCL5 (**Figure 3D**).

In the context of pro-inflammatory cytokines, we observed a concentration-dependent increase in IL-6 for cannabis smoke exposure at 10% (*p*<0.05 – **Figure 3F**) and a trend for IL-8 (*p*>0.05 – **Figure 3E**). In contrast, tobacco smoke displayed a bell-shaped concentration-response curve with a concentration-dependent increase observed from 0.625% to 5% with a decrease observed at concentrations beyond 10%.

### Analysis of formoterol/budesonide treatment on cannabis smoke exposure-induced alterations in epithelial cell viability, barrier function, and immune profile

The common anti-inflammatory formoterol/budesonide was examined in the context of cannabis exposure using 10% smoke conditioned media as a concentration that maximized a combination of cell viability and impact on barrier function and immune profile (**Figure 4**). A single concentration of formoterol/budesonide (10nM/100nM) was used as this concentration is capable of displaying anti-inflammatory responses in airway epithelial cells (50, 52).

**Figure 4.**
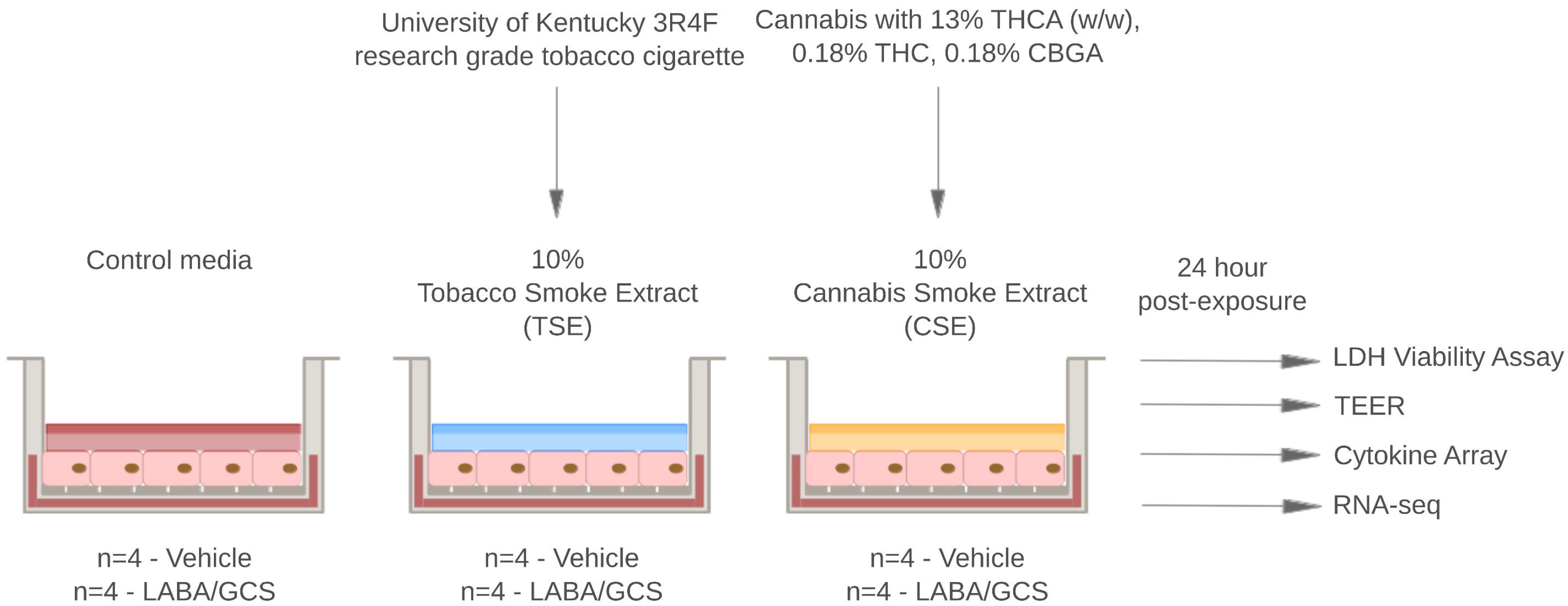
Formoterol/Budesonide Intervention Experiment Work-Flow. Analysis of formoterol/budesonide treatment on cannabis smoke exposure-induced alterations in epithelial cell viability, barrier function, and immune profile relative to tobacco smoke. A single concentration of the anti-inflammatory formoterol/budesonide (10nM/100nM) was examined in the context of cannabis exposure using 10% smoke conditioned media. Calu-3 cells were exposed to either control, TSE, or CSE with either vehicle or formoterol/budesonide (n=4) for 24 hours before downstream analysis via LDH viability assay, TEER, cytokine array, and RNA-seq.

### Formoterol/budesonide combination treatment does not impact epithelial cell viability while preventing cannabis induced decreases in barrier function

The 10% dilution of cannabis failed to impact cell viability in the concentration-response study (**Figure 2A**), which was confirmed in the presence or absence of formoterol/budesonide (**Figure 5A**). Similarly, tobacco smoke exposure in the presence or absence of formoterol/budesonide failed to impact cell viability. The 10% dilution of cannabis reduced epithelial cell barrier function in the concentration-response study (p<0.05 - **Figure 2B**). The formoterol/budesonide intervention study confirmed that 10% cannabis smoke exposure reduced epithelial cell barrier function and that this change was minimized by formoterol/budesonide intervention (*p*<0.05 – **Figure 5B**). Similar results were observed for tobacco smoke. Importantly, formoterol/budesonide intervention increased barrier function measurements in control media exposed epithelial cells in the absence of any smoke conditioned media.

**Figure 5.**
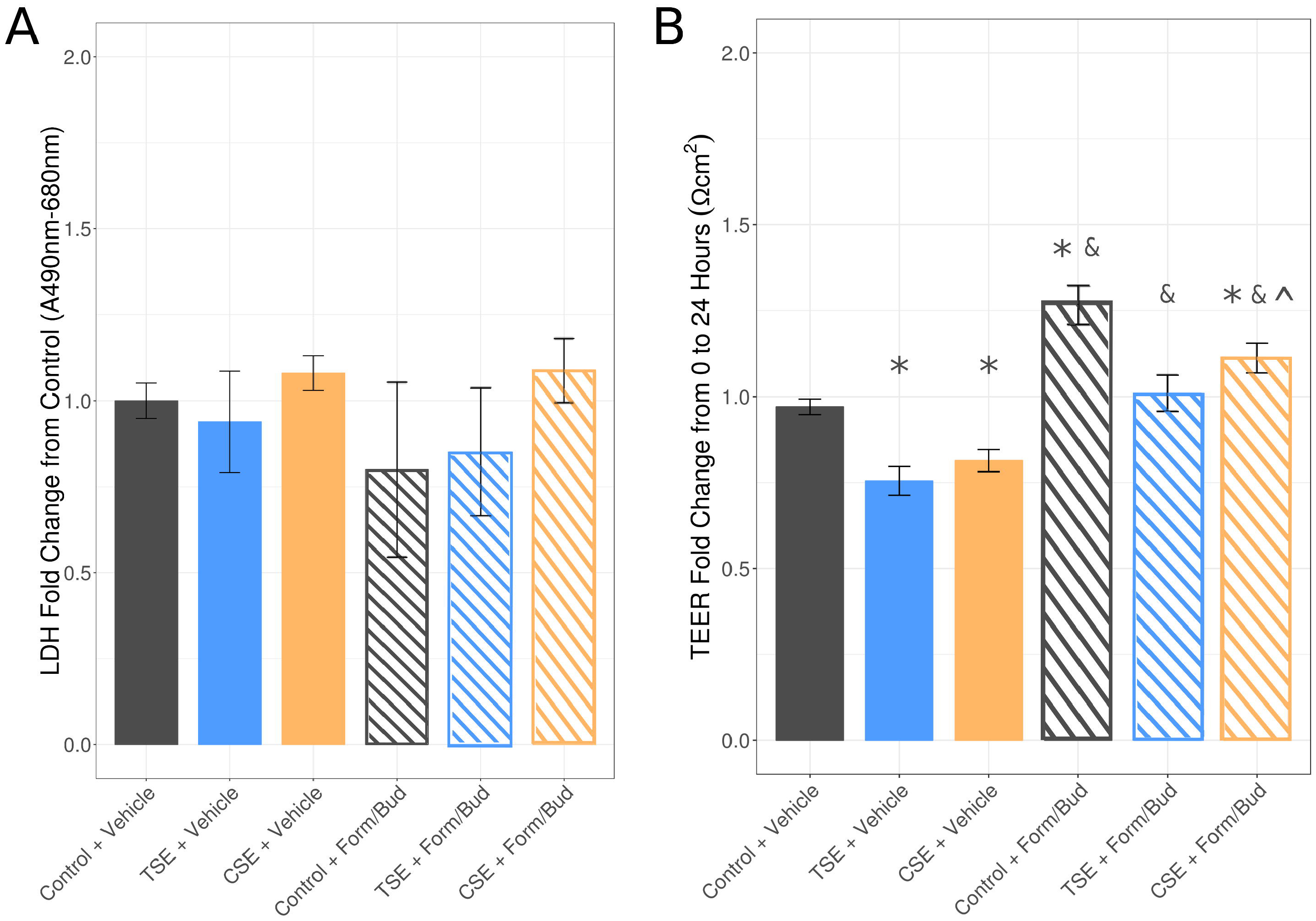
Effect of cannabis smoke exposure and formoterol/budesonide intervention on airway epithelial cell viability and barrier function. Calu-3 cells were exposed to 10% CSE (orange) or TSE (blue) for 24 hours. ***A*)** Cell viability was assessed via LDH assay where each exposure was calculated as a fold change from untreated cells. ***B*)** TEER was measured at 0 and 24 hours post-exposure and fold change for each exposure was calculated and compared to untreated cells. * = p<0.05 relative to control+vehicle, & = <0.05 relative to corresponding control, ^ = p<0.05 relative to TSE+formoterol/budesonide - Tukey HSD.

### Formoterol/budesonide combination treatment selectively modulates epithelial cell immune profile in response to cannabis exposure

In the context of epithelial cell barrier repair mediators, formoterol/budesonide intervention did not impact increases in TGF-α and PDGF-AA induced by 10% cannabis smoke conditioned media (*p*<0.05 – **Figure 6A-B**).

**Figure 6.**
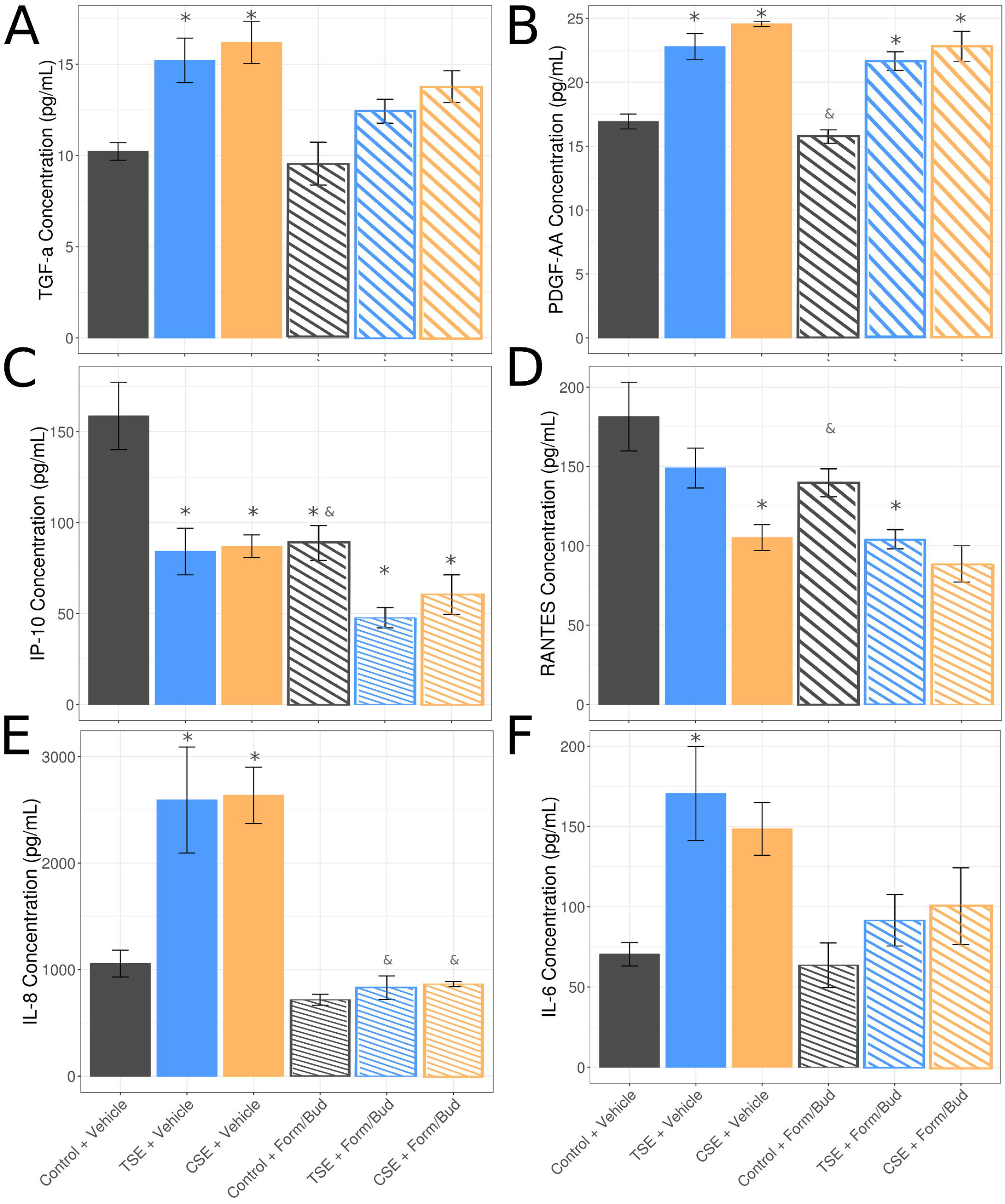
Effect of cannabis smoke exposure and formoterol/budesonide intervention on select cytokine production. The same 6 cytokines selected during the dose-response experiment were examined to determine the effects formoterol/budesonide intervention has on 10% smoke extract exposure. ***A*)** TGF-α, ***B*)** PDGF-AA, ***C*)** IP-10, ***D*)** RANTES, ***E*)** IL-8 and ***F*)** IL-6. * = p<0.05 relative to control+vehicle, & = <0.05 relative to corresponding control, Tukey HSD.

In the context of antiviral immune mediators, formoterol/budesonide intervention potentiated the reduction in IP-10 levels induced by 10% cannabis smoke conditioned media (*p*<0.05 – **Figure 6C**), with a smaller trend observed for RANTES/CXCL5 (*p*<0.05 – **Figure 6D**).

In the context of pro-inflammatory cytokines, formoterol/budesonide intervention was able to completely normalize increases in IL-8 (*p*<0.05 – **Figure 6E**) and to a lesser extent IL-6 (*p*<0.05 – **Figure 6F**) that are induced by 10% cannabis smoke conditioned media.

For each of the above cytokines, similar results were observed for 10% tobacco smoke conditioned media exposure.

### Transcriptomic analysis of airway epithelial cells exposed to cannabis and tobacco smoke, and formoterol/budesonide intervention

The analysis of epithelial cell viability, barrier function, and immune profile in the presence or absence of formoterol/budesonide (**Figures 1-6**) revealed that similar impacts were induced in airway epithelial cells by cannabis and tobacco smoke exposure. We therefore explored the similarities and differences between cannabis and tobacco smoke exposure at a transcriptional level. A parallel analysis was performed to determine if the impact of formoterol/budesonide interventions were shared between cannabis and tobacco smoke exposed airway epithelial cells.

A global transcriptomic analysis of airway epithelial cells from the smoke and formoterol/budesonide intervention study was performed using RNA-sequencing. A total of 300 and 598 genes were significantly up-regulated in TSE and CSE (q < 0.05), respectively (**Supplemental Tables 2, 4**), and 279 and 623 genes were significantly down-regulated in TSE and CSE (q < 0.05) (**Supplemental Tables 3, 5**). Formoterol/budesonide intervention resulted in an additional shift in gene expression with 1316 differentially expressed genes in CSE+drug versus CSE alone (**Supplemental Table 6**), and 333 differentially expressed genes in TSE versus TSE alone (**Supplemental Table 7**).

Before analyzing the nature of differentially expressed genes, we first clustered all samples (including replicates) based on their transcriptomic profiles and visualized their overall similarities in gene expression via non-linear multi-dimensional scaling (NMDS) using the “vegan” package in R (v.3.4.3). In the NMDS plot, the distance between samples reflects similarity in gene expression profiles (**Figure 7**). The transcriptomic profiles clustered distinctly into four main groups according to condition: Control+Vehicle, Control+Intervention Smoke(cannabis or tobacco) +Vehicle, and Smoke (cannabis or tobacco) +Intervention. Further, the transcriptomes of cannabis and tobacco smoke exposed cells clustered closely to one another but also formed distinct groups by smoke type. The smoke treated samples also clustered distinctly in the formoterol/budesonide-treated samples, although drug intervention resulted in an additional shift in gene expression profiles. According to NMDS clustering, this shift appears to occur along a similar directional axis as that for formoterol/budesonide-treated control samples (p < 0.001), which may reflect the addition of an expression signature unique to formoterol/budesonide (**Figure 7**). Differentially expressed genes and pathways in each condition are further analyzed in detail below.

**Figure 7.**
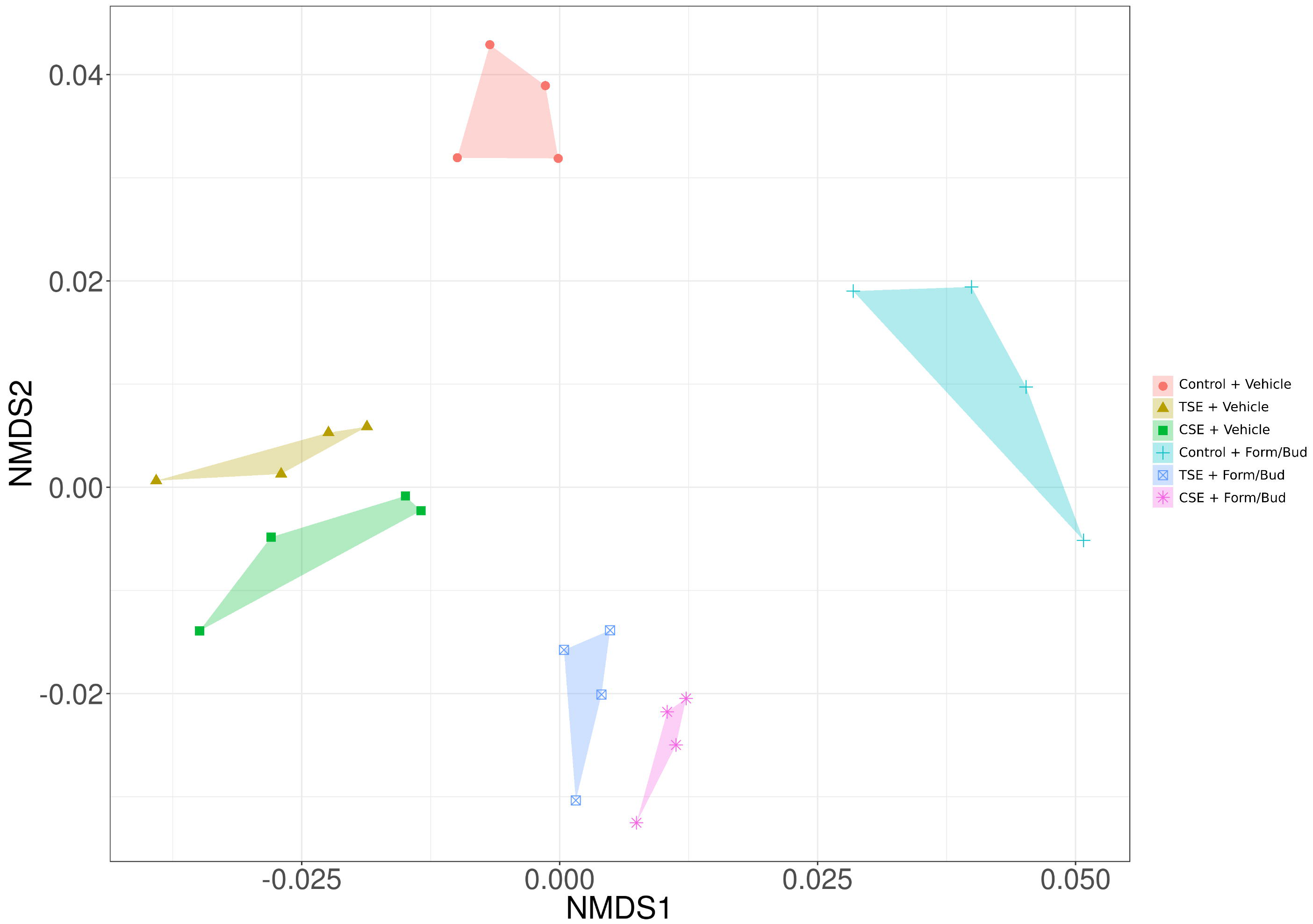
Effect of cannabis smoke exposure and formoterol/budesonide intervention on airway epithelial transcriptional profiles. Non-linear multi-dimensional scaling was performed to assess the overall similarity between cannabis and tobacco exposure on global transcriptomic profiles. Both cannabis and tobacco smoke exposure result in significant differentiation from control in terms of the overall transcriptional profile (ANOSIM, *p* < 0.004, *p* < 0.02, respectively). After intervention with formoterol/budesonide, a strong intervention signature is observed (ordination vector, *p* < 0.001). Cannabis and tobacco exposure groups significantly vary from control following treatment with formoterol/budesonide (ANOSIM, *p* < 0.009, *p* < 0.03, respectively) and the intervention does not return the smoke profiles to that of the control. Cannabis and tobacco profiles do not overlap, implying subtle differences between the two smokes however, this was not found to be significant.

### Cannabis vs tobacco smoke exposure produces a correlated shift in gene expression

As demonstrated in the NMDS plot, both cannabis and tobacco smoke exposure resulted in a similar shift in gene expression that was distinguishable from control media exposed cells (ANOSIM, *p* < 0.004, *p* < 0.02, respectively - **Figure 7**). In addition, because CSE and TSE samples clustered closely by NMDS, this suggests that the transcriptomic response to both smokes may be correlated. The transcriptomes of cannabis and tobacco smoke exposed cells clustered closely to one another but also formed distinct groups by smoke type. This suggests that there is a large shared component in the transcriptomic response to smoke with subtle differences due to the nature of the smoke extract exposure.

To examine the similarity in gene expression response to CSE versus TSE in more detail, we compared the log_2_ fold changes of all genes following TSE versus CSE exposure (**Table 1**), and evaluated the overlap in differentially expressed genes in both conditions (**Figure 8**). The gene expression response to TSE versus CSE exposed cells was highly correlated (r = 0.695, *p*<1×10^−15^). In addition, there was significant overlap in differentially expressed genes following cannabis and tobacco exposure, with 391 genes in common (*p*<1×10^−15^ - **Figure 8**– Red and purple circle overlap in Venn Diagram). Direct comparison of the CSE-induced versus TSE-induced transcriptomes revealed only seven genes that were differentially expressed between the two smoke types (**Table 2 -**Orange points in **Figure 8**), again indicating considerable similarity in transcriptomic profiles between both cannabis and tobacco smoke exposures (**Table 2**). Despite there being only seven differentially expressed genes between CSE and TSE exposure, these genes may be of interest in the context of airway epithelial cell biology following smoke exposure. For example, TNFRSF10A (Death Receptor 4, DR4) exhibited significantly lower expression in CSE versus TSE exposed cells, which suggests that CSE-exposed cells may be under-sensitized to TRAIL-mediated apoptosis. In addition, *SQSTM1/p62* (Sequestosome 1) was up-regulated in TSE and down-regulated in CSE exposed cells (**Figure 8**). Elevated expression of this gene has been associated with aggressive tumor behavior in early-stage non-small cell lung cancer(53).

**Figure 8.**
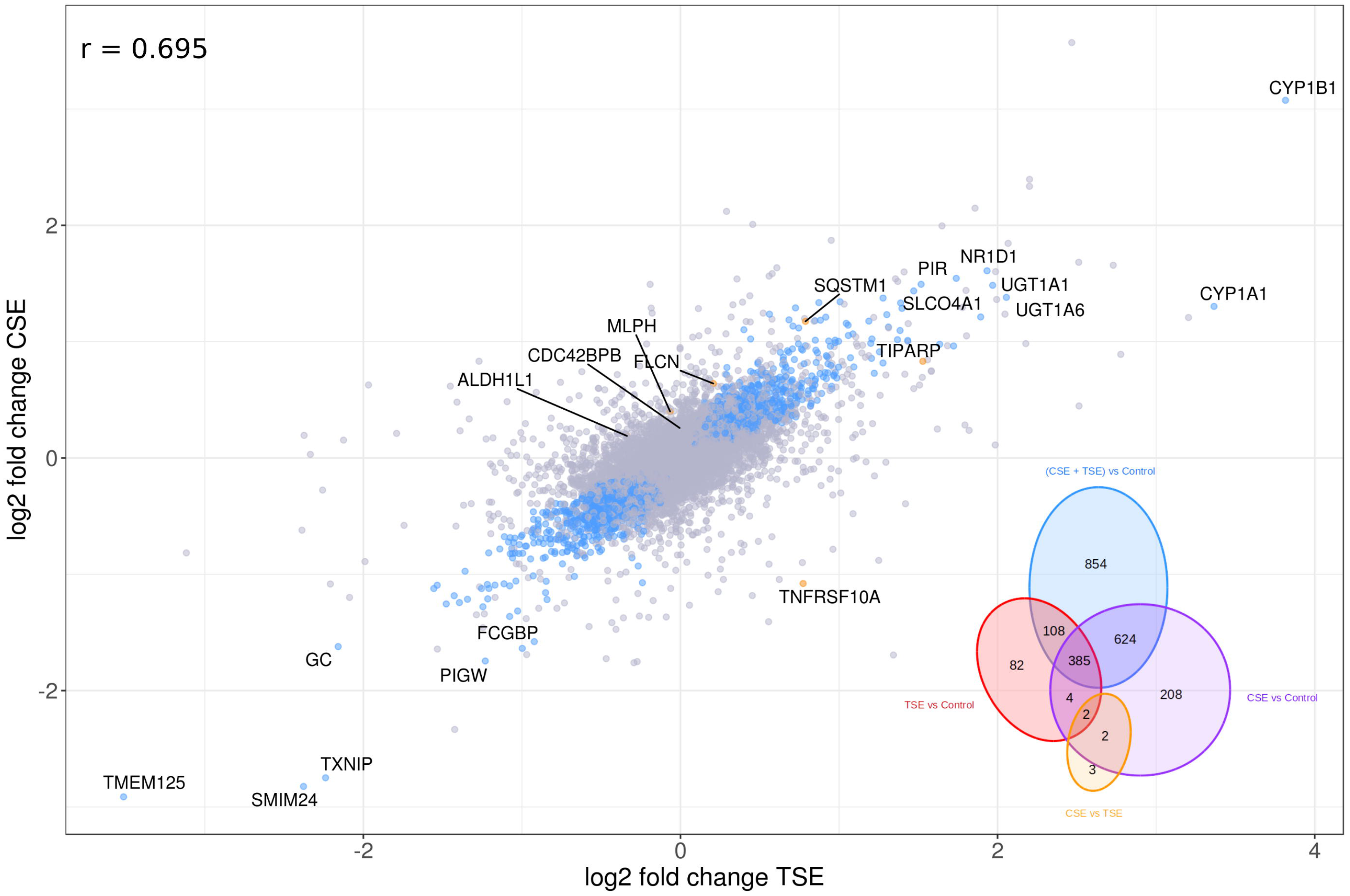
Comparison of cannabis and tobacco gene expression profiles. The effects of cannabis and tobacco smoke exposure on airway epithelial gene expression were compared by plotting log_2_ fold change of TSE/Control against log_2_ fold change of CSE/Control. High positive correlation is seen between the two smokes (Pearson correlation, r = 0.695, *p* < 1×10^−15^). 911 genes showed significant (*q* < 0.05) expression patterns common to both cannabis and tobacco (as determined by analysis of differential expression between (CSE + TSE) vs Control) and are highlighted in blue. Only 7 genes, highlighted in orange), were determined to be differentially expressed (*q* < 0.05) between cannabis and tobacco. A Venn diagram was used to show the break-down of differentially expressed genes between comparisons.

### Impact of formoterol/budesonide treatment on global gene expression

Formoterol/budesonide treatment induced a shift in gene expression in control (non-smoke) cells that deviated from vehicle exposed control (non-smoke) cells (**Figure 7**). The direction of the transcriptomic shift was paralleled in formoterol/budesonide treated cannabis-exposed and tobacco-exposed cells (*p*<1×10^−5^, **Figure 7**), and again samples from both smoke types clustered distinctly in the NMDS plot, suggesting subtle differences due to the nature of the smoke extract exposure in the context of formoterol/budesonide treatment. Given that formoterol/budesonide treatment did not fully normalize the gene expression signature by overlapping with control samples, our data suggests that there are smoke-induced transcriptomic effects that remain in cannabis and tobacco exposed samples after drug intervention.

### Analysis of differentially expressed genes following cannabis vs tobacco smoke

Given the considerable correlation in gene expression response to cannabis and tobacco smoke, we pooled the transcriptomes of CSE and TSE exposed cells together to more sensitively detect the full set of smoke-induced differentially expressed genes. After pooling, we detected 911 differentially expressed genes by TSE+CSE smoke (**Supplemental Tables 5-6**), which are visualized as blue points in **Figure 8.**

The most dramatically up-regulated genes following TSE and CSE exposure are the cytochrome P450 genes, *CYP1A1* and *CYP1B1* (**Figure 8**), which are 1.3-3.4-and 3.1-3.8 fold up-regulated by TSE and CSE smoke, respectively. This is consistent with previous literature, since these genes are known to be up-regulated in smoke exposed lung tissue(54), and are among the most highly induced genes in cells exposed to the smoke-associated carcinogen, benzo-a-pyrene(55, 56), which is present in both tobacco and cannabis smoke tar.

To test this idea further, we examined other genes known to be induced by benzo-a-pyrene based on previous microarray or RNA-seq studies(55–57). In addition to *CYP1A1* and *CYP1B1*, key genes that have been implicated in the benzo-a-pyrene response include *AP1* (Fos, Jun), *MYC*, *CDH1*, *TUBB5*, *APC*, *CAV*, *TP53*, *PCNA*, *STAT3*, *ERBB2*, *E2F1*, *NQO1*, *NR2F*, *EPHX1*, and NF-κB and Wnt/ß-catenin (*CTNNB1*) pathways. Of these genes, 6 were significantly differentially expressed in both TSE and CSE treatments in our study (**Supplemental Tables 8-9**).

### Function enrichment analysis detects increased oxidative stress and oncogenic pathways and lowered antiviral responses in smoke-exposed cells

To assess the transcriptomic response to smoke at the level of biological pathways and functions, we performed function enrichment analysis using several functional annotation ontologies (**Figure 9**). The top GO (Gene Ontology) enrichment among genes up-regulated by smoke exposure was “cellular response to oxidative stress” (p <0.00004). This may be a direct result of the dramatic up-regulation in CYP1A1 and CYP1B1 expression, since elevated CYP expression is known to cause oxidative stress to cells (58). Consistent with this finding, the top enriched pathway among smoke-induced genes was “Oxidative Stress Induced Gene Expression Via Nrf2”. The Nrf2 gene (*NFE2L2*) itself was significantly up-regulated in both TSE and CSE treatments, and the Nrf2 pathway has been previously identified as an important regulatory response to benzo-a-pyrne exposure(59). Other oxidative stress related genes up-regulated in both conditions include *NQO1, GSTP1*, and *OSGIN1*.

**Figure 9.**
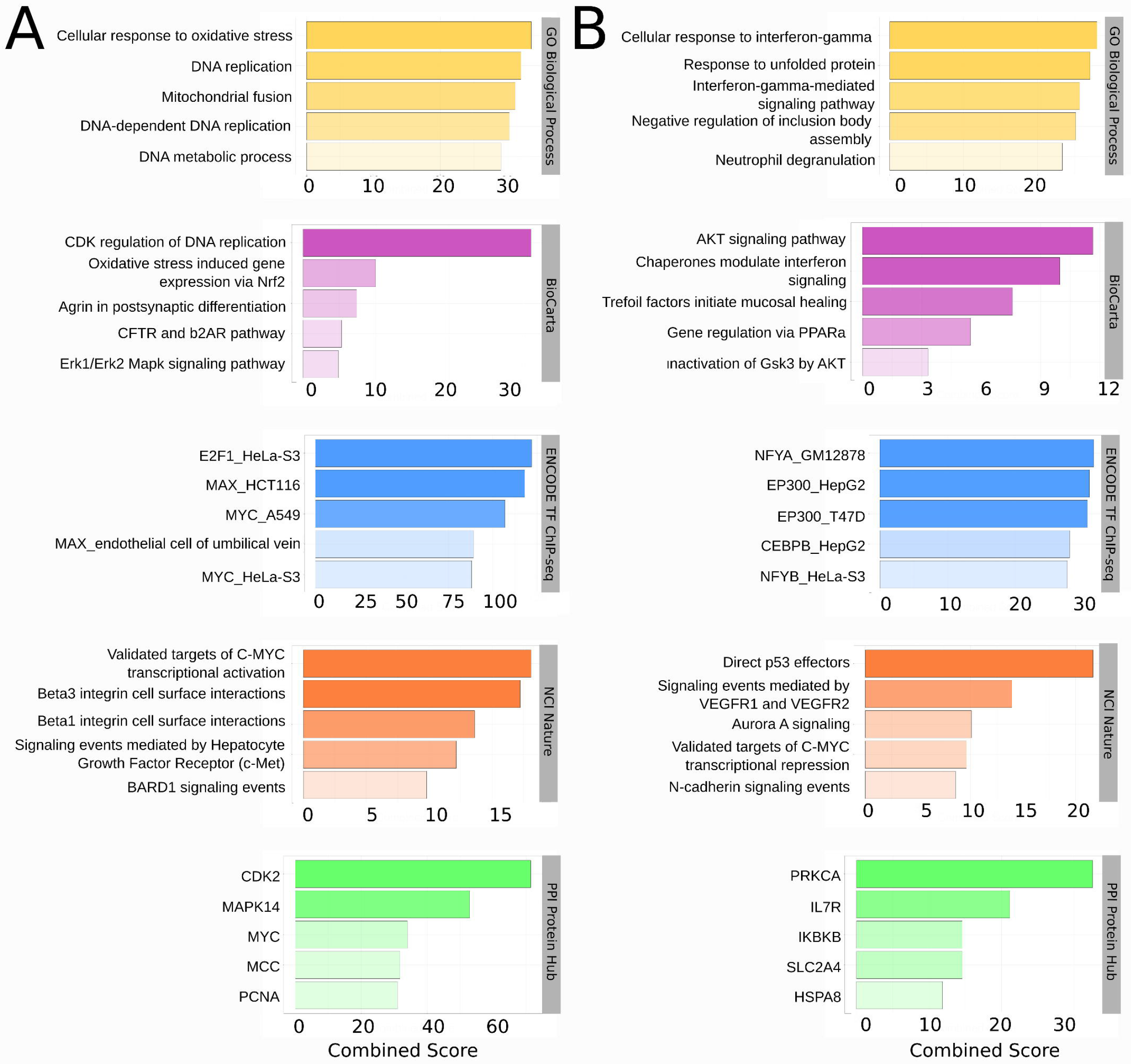
Functional enrichment of genes common between cannabis and tobacco. The 911 genes identified as differentially expressed in cannabis and tobacco compared to control were separated into ***A*)** up- and ***B*)** down-regulated lists and submitted to EnrichR for functional enrichment analysis. The top 5 terms (as determined by EnrichR Combined Score) were plotted for selected ontologies: GO Biological Process (yellow), BioCarta Pathway (purple), ENCODE Transcription Factor (TF) ChIP-seq Data (blue), NCI Nature (orange), and PPI Protein Hubs (green).

GO enrichment analysis of smoke-induced genes also detected an over-representation of genes involved in DNA replication (**Figure 9A**), which likely reflects the regulatory control of cell proliferation and DNA damage response during chemical and/or oxidative stress that has also been observed in previous benzo-a-pyrene studies(55–57). Analysis of enriched pathways and protein-protein interaction (PPI) hub proteins suggests that gene expression changes related to these functions may occur in part via CDK2 interactions (**Figure 9**).

Genes up-regulated by smoke also overlapped significantly with previous cancer gene expression signatures. The oncogene *MYC* was significantly up-regulated in both CSE and TSE exposed cells, and Myc targets and PPI partners were all significantly enriched among the smoke-induced genes. In addition to elevated expression of *MYC*, which is commonly observed in cancer transcriptomes, we also observed enrichment of the P53 pathway in both smoke exposures. “Direct P53 effectors” (NCI-Nature Pathways Database) was the top enriched term among genes down-regulated by smoke exposure. Down-regulation of the P53 pathway is also an expression signature in some cancers where it may contribute to resistance to apoptosis.

Finally, in agreement with earlier cytokine analysis results suggesting a dampened immune response to pathogens in smoke-exposed cells, the top enriched GO term for down-regulated genes (**Figure 9B**) was “cellular response to interferon-gamma”. To further test the idea that smoke exposure may inhibit IFN-mediated immune responses, we examined 10 targets of IFN-mediated antiviral pathway (see Methods). Six of these (*MX1, ISG15, IFITM3, RSAD2, MDA5, IFNB1*) show significantly reduced gene expression levels in smoke-exposed cells.

### Impact of formoterol/budesonide treatment on global gene expression

Next, we sought to examine the transcriptomic response of smoke-exposed cells to formoterol/budesonide treatment, with the goal of exploring: 1) differences in drug-response between CSE and TSE exposed cells; and 2) impact of drug treatment on the smoke-induced gene expression changes described earlier.

Formoterol/budesonide treatment induced a shift in gene expression in control (non-smoke) cells that deviated from vehicle exposed control (non-smoke) cells (**Figure 7**). The direction of the transcriptomic shift was paralleled in formoterol/budesonide treated cannabis-exposed and tobacco-exposed cells (*p*<1×10^−5^, **Figure 7**), and again samples from both smoke types clustered distinctly in the NMDS plot, suggesting subtle differences due to the nature of the smoke extract exposure in the context of formoterol/budesonide treatment. Given that formoterol/budesonide treatment did not fully normalize the gene expression signature by overlapping with control samples, our data suggests that there are smoke-induced transcriptomic effects that remain in cannabis and tobacco exposed samples after drug intervention.

As before with control treated cells, CSE and TSE exposure in formoterol/budesonide-treated cells induced a highly correlated transcriptomic response (**Supplementary Figure 1**). To identify unique expression patterns in formoterol/budesonide and smoke-exposed cells versus only smoke-exposed cells, we directly compared these datasets to identify differentially expressed genes and visualized their log_2_ fold changes relative to controls (**Figure 10**). As expected, the most dramatically up-regulated gene that was exclusive to formoterol/budesonide+smoke exposed cells was HSD11B2 (Corticosteroid 11-β-dehydrogenase). The expression of *NEU1*, whose up-regulation is associated with idiopathic pulmonary fibrosis(60), was also significantly reduced in formoterol/budesonide-treated control cells, but remained at higher relative expression with smoke exposure (**Figure 11**). Finally, key expression patterns associated with oncogenesis, such as elevated *MYC* levels, were dampened in formoterol/budesonide-exposed cells (**Figure 11**).

**Figure 10.**
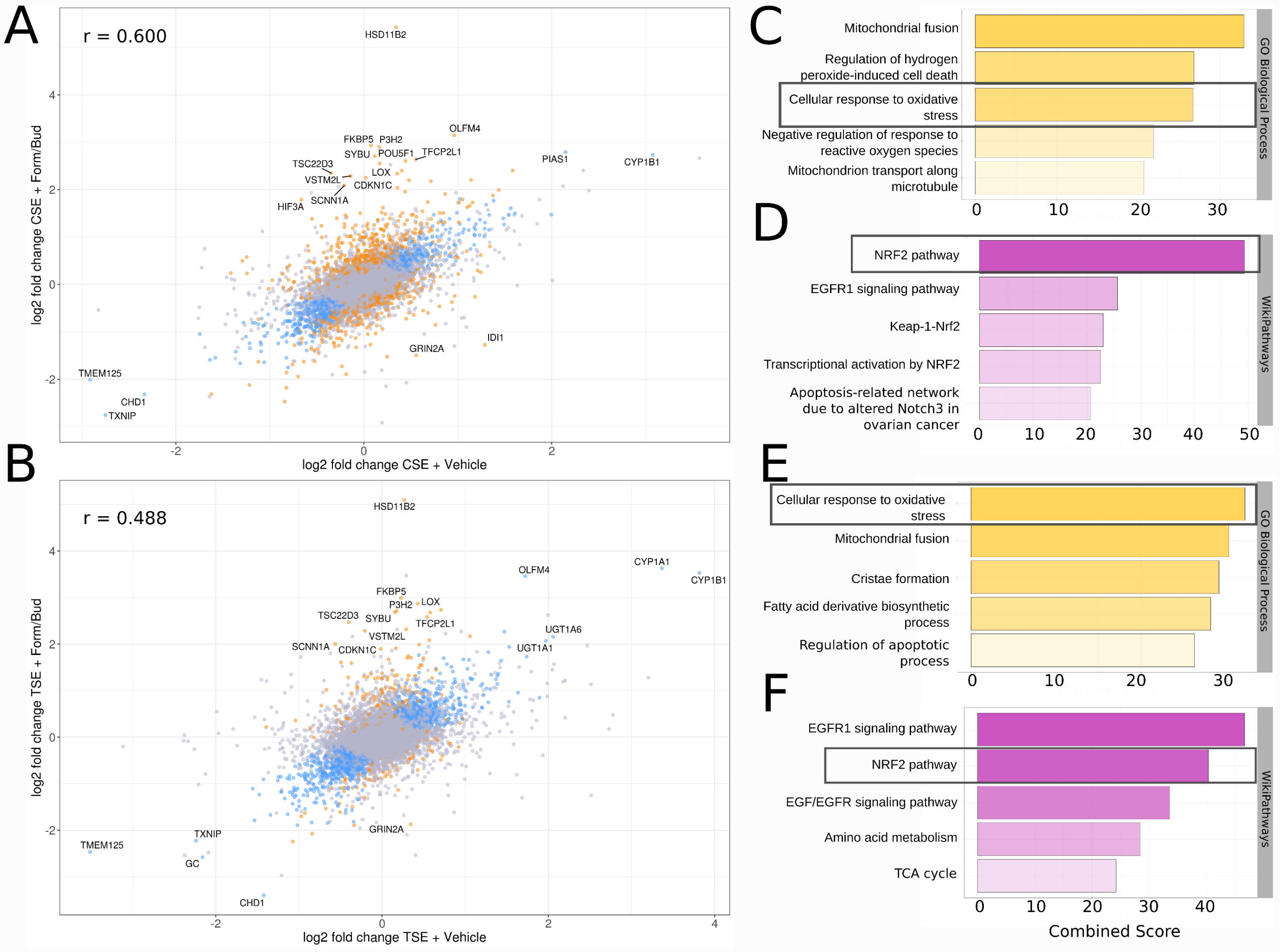
Effect of formoterol/budesonide intervention on cannabis- and tobacco-induced gene expression profile changes. The effect of formoterol/budesonide intervention on airway epithelial gene expression was assessed in the context of ***A)*** cannabis and ***B*)** tobacco smoke exposure. ***A*)** Log_2_ fold change of CSE/Control against log_2_ fold change of CSE + formoterol/budesonide. ***B*)** Log_2_ fold change of TSE/Control against log_2_ fold change of TSE + formoterol/budesonide. Genes showed to be significantly differentially expressed (*q* < 0.05) in (smoke + (smoke + formoterol/budesonide)) vs control are highlighted in blue. Formoterol/budesonide specific genes shown to be significantly differentially expressed between smoke vs (smoke + formoterol/budesonide) are highlighted in orange. **C-D)** GO Biologics and WikiPathways gene ontologies for log_2_ fold change of CSE + formoterol/budesonide against log_2_ fold change of CSE/Control. **E-F)** GO Biologics and Wikipathways gene ontologies for log_2_ fold change of TSE + formoterol/budesonide against log_2_ fold change of TSE/Control

**Figure 11.**
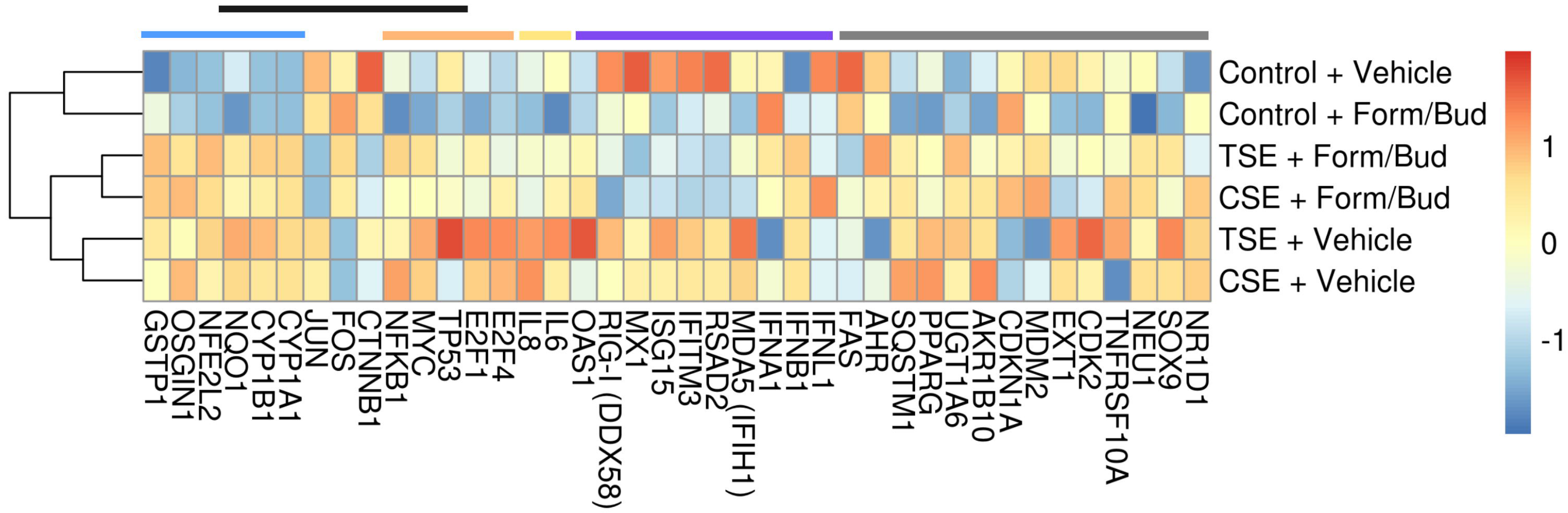
Effect of formoterol/budesonide intervention on top ontology drivers. Genes of interest were selected from literature based on or observations of increased oxidative stress, benzo-a-pyrene-related pathway enrichment, and a dampening of antiviral responses, to assess the effects of formoterol/budesonide on these effects. The impact of formoterol/budesonide intervention was examined for genes involved oxidative stress (blue bar – top of heat map), benzo-a-pyrene related genes (black bar – top of heat map), oncogenic (orange bar – top of heat map), antiviral genes (purple bar– top of heat map), markers of inflammation (yellow bar – top of heat map) and other (grey bar – top of heat map).

Next, we performed function enrichment analysis for genes that were differentially expressed in the formoterol/budesonide and smoke-exposed cells, and did so separately for CSE and TSE samples. Interestingly, as before with the control samples, the top function enrichments for up-regulated genes in both CSE and TSE relate to oxidative stress pathways (**Figure 10**). Consistent with this finding, the dramatic up-regulation of *CYP1A1* and *CYP1B1* genes remains as an expression signature of formoterol/budesonide and smoke-exposed cells for both smoke types (**Figure 10** and **Supplementary Figure 1**). As shown in the expression heat map for select genes (**Figure 11**), these and other key oxidative stress response genes (e.g., *NQO1, NFE2L2,* and *OSGIN1*) are relatively unaffected by drug treatment and remain at high relative expression compared to control samples (**Figure 11**). Also as expected, genes involved in pro-inflammatory and antiviral responses were also down-regulated (e.g. *CXCL8/IL-8, RSAD2*, and *ISG15*, **Figure 11**) in formoterol/budesonide + smoke exposed cells compared to smoke-only cells, consistent with the cellular pathways known to be targeted by LABA/GCS therapy

Thus, we conclude that combination formoterol/budesonide therapy had little to no effect on the smoke-related induction of oxidative stress pathways, but resulted in reduced expression levels of pro-inflammatory and antiviral pathways.

### The impact of cannabis smoke exposure on markers of oxidative stress in airway epithelial cells

Differential gene expression analysis revealed that both cannabis and tobacco smoke exposure resulted in upregulation of CYP expression, and associated oxidative stress pathways that may be a direct consequence of elevated CYP activity (p<0.05 - **Figure 12A-B**) that remained significantly elevated following LABA/GCS treatment.

**Figure 12.**
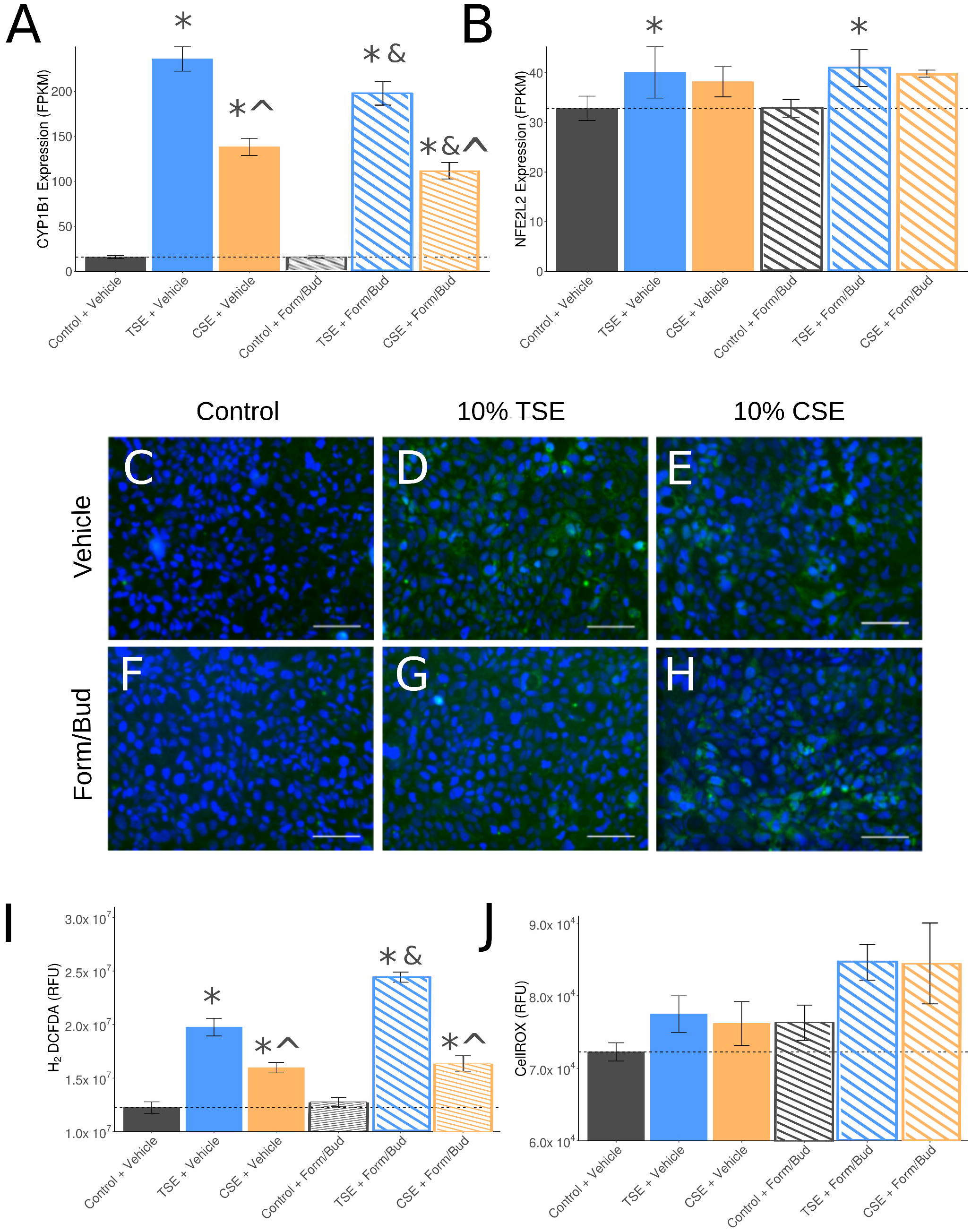
Effect of cannabis smoke exposure and formoterol/budesonide intervention on airway epithelial cell oxidative stress. Since previous results show that both cannabis and tobacco increase airway epithelial oxidative stress, an effect unable to be attenuated by formoterol/budesonide intervention, *in vitro* cell microscopy was performed to validate. Expression of the oxidative stress-related genes ***A***) *CYP1B1* and ***B***) *NFE2L2* were quantified using RNA-seq data and correlated to in vitro experiments with the oxidative stress cell permeable dyes, H_2_DCFDA (**C-E**) and CellROX (**F-H**) following cannabis or tobacco exposure in the presence or absence of formoterol/budesonide and quantified (**I-J**). * = p<0.05 relative to control+vehicle, & = <0.05 relative to corresponding control, ^ = p<0.05 relative to TSE - Tukey HSD.

Complementary imaging and quantification experiments using reactive oxygen species-sensitive fluorescent dyes in airway epithelial cells exposed to cannabis smoke in the presence or absence of formoterol/budesonide confirmed gene expression analysis (**Figure 12C-H)**. Quantification of staining intensity revealed increased reactive oxygen species generation in airway epithelial cells exposed to 10% cannabis smoke conditioned media as measured by H_2_DCFDA, with similar trends observed with CellROX. Similarly, tobacco smoke exposure resulted in an increase in both signals. Formoterol/budesonide intervention potentiated the reactive oxygen species signal in both cannabis and tobacco exposed cells.

## DISCUSSION

Global recreational cannabis use is a potentially important public health issue that would benefit from experimental evidence to inform policy, regulations, and individual user practices. Comparative analyses between cannabis and tobacco smoke (25, 26), the latter long reported to have negative impacts on respiratory health (4–19), may help provide context and provide clinically relevant evidence. To address this unmet need and leverage an extensive history of *in vitro* tobacco smoke exposure research, we performed a comparative study between cannabis and tobacco smoke exposure in the Calu-3 human airway epithelial cells using concentration-response and pharmacological intervention study designs. Our results show striking similarities between cannabis and tobacco smoke exposure on barrier function, suppression of antiviral mediators, potentiation of pro-inflammatory mediators, and induction of oncogenic and oxidative stress gene expression signatures. Furthermore, we demonstrate that a common class of anti-inflammatories, LABA/GCS, were unable to prevent cannabis smoke-induced reductions in antiviral pathways or normalize induction of oncogenic and oxidative stress responses. Collectively our data suggest that cannabis smoke exposure is not innocuous and may possess many of the deleterious properties of tobacco smoke.

The function of the airway epithelium is to provide the lung a mechanical and immunological barrier to the outside world and protect the world from inhaled insults including air pollution, bacteria, and viruses (35, 36). Any perturbation in the airway epithelium may lead to host susceptibility to infection and subsequent lung damage that if not controlled effectively, could manifest in lung pathology. Indeed, chronic airway diseases including asthma(61), chronic obstructive pulmonary disease(62), and idiopathic pulmonary fibrosis (63), are associated with abnormalities in airway epithelial cell biology. Our data demonstrate that cannabis smoke exposure is able to induce mild impacts on barrier function, measured by TEER, without impacting cell viability. The mechanism(s) by which TEER is reduced by cannabis were not determined in our study, but absence any changes in cell viability it is possible that cell-cell junctions could have been disrupted, as has been reported with tobacco smoke (64). Transwell membrane permeability experiments with fluorescently labeled substrates would help define the degree that the airway epithelium barrier function is compromised following cannabis smoke exposure. Irrespective of the mechanism, reduced cell-cell junctions as indicated by reduced N-cadherin signaling events in our ontology analyses, may result from cellular differentiation processes regulating repair mechanisms(65). In response to cannabis smoke exposure, beta-catenin shuttling from E-cadherin membrane junction sites to transcriptional locations in the nucleus may occur to facilitate gene expression associated with repair. Importantly, aberrant beta-catenin signaling cells is associated with oncogenic gene expression signatures and cancer development(66), which could be important in the context of lung health in the cannabis smoker. Independent of aberrant transcriptional regulation resulting from disrupted epithelial barrier function, the reduced mechanical impedance offers an easier access to the lung for opportunistic pathogens.

To complement the mechanical barrier of the lung, the airway epithelium is able to produce and host-defence peptides and antiviral mediators to protect from bacteria and viruses, respectively(35, 36). Tobacco smoke exposure has been reported to compromise the ability of airway epithelial cells to effectively control both bacterial and viral insults (4–6, 9–12). Our data demonstrating striking similarities in the epithelial immune profile in response to cannabis and tobacco smoke exposure, suggesting that the former will also impact bacterial and viral host defences. Tobacco smoke has been demonstrated to impact host defence peptide induction in airway epithelial cells by nontypeable *Haemophilus influenzae* with a concomitant increase in IL-8 expression (4). Unfortunately, our experimental dataset did not include a pathogen challenge, precluding our ability confirm the tobacco smoke induced suppression of host defence peptides or to extend the results to cannabis exposure. In the context of viral exposures, type I interferons (IFNs) are capable of rapidly inducing interferon stimulated genes (ISGs) through the type I IFN receptor to help tackle various components of virus replication, assembly and budding(67). Tobacco smoke exposure has been shown to impact antiviral immunity in airway epithelial cells in response to human rhinovirus-16, with a reduction in IP-10 and RANTES(6). The tobacco-induced reduction in IP-10 and RANTES was associated with greater rhinovirus production. Tobacco smoke has also been demonstrated to impair airway epithelial cell antiviral immunity mediated by IFN-γ in response to respiratory syncytial virus exposure(68), which in turn could impact IP-10 and RANTES production(69). Our cytokine and transcriptomic data confirm the tobacco smoke impairment of IP-10 and RANTES and IFN-γ ontologies and show that this is conserved for cannabis smoke exposure. We further demonstrate using our transcriptomics dataset that a diverse selection of ISGs were also attenuated with both cannabis and tobacco smoke exposure, consolidating a common phenotype between both smoke exposures. Collectively, although our experimental designs lack the mechanistic linkage between cannabis smoke exposure and increased susceptibility to viral or bacterial infections, our data strongly mirror those for tobacco smoke exposure, which has been mechanistically linked to compromised host immunity to pathogens.

Chronic bronchitis is a key pathological feature of chronic obstructive pulmonary disease (70). Clinical management of chronic bronchitis may include LABA/GCS interventions including formoterol/budesonide combination(71–73). LABA/GCS are transcriptionally active in airway epithelial cells resulting in broad anti-inflammatory activities (49). Despite their widespread use, recent analyses of LABA/GCS efficacy and mechanisms have demonstrated that these compounds may suppress antiviral immunity, which could increase susceptibility to subsequent bacterial infections (12, 71–74). Importantly, the efficacy of LABA/GCS therapies may be compromised by tobacco smoke exposure(8, 13), which could be contributing to the reduced efficacy in certain patient populations. As cannabis smoke exposure is also associated with development of chronic bronchitis and may be treated with LABA/GCS in the clinic, we investigated the responses of airway epithelial cells to a commonly prescribed formulation of formoterol/budesonide. Our LABA/GCS intervention data confirm our concentration-response study that observed a cannabis-induced reduction in IP-10 and RANTES. We further demonstrate that LABA/GCS intervention augments the cannabis and tobacco smoke-induced reduction of antiviral mediators at both the protein (IP-10 and RANTES) and gene expression (ISGs – *VIPERIN/RSAD2, OAS1, MDA5, RIG-I*) level. The implications of our data suggest that prescription of LABA/GCS for cannabis smokers should be used with caution as this may impair antiviral responses and predispose individuals to bacterial infections.

Extensive evidence exists that tobacco smoke and biomass exposure, including dried wood, animal dung, or charcoal, are risk factors for the development of chronic bronchitis, emphysema, and lung cancers (75–78). In contrast, the existing evidence suggests that chronic cannabis smoke exposure results in only a chronic bronchitis phenotype with little evidence of emphysema (25–28). Furthermore, unlike tobacco and biomass exposure, which are accompanied by an dose-dependent risk for development of lung cancers, a similar relationship has not been observed for chronic cannabis users despite the presence of carcinogens in cannabis smoke(29). Of particular note, we observed that cannabis smoke significantly up-regulated the proto-oncogene *MYC,* which has previously been observed in airway epithelial cells exposed to tobacco smoke (79) or benzo-a-pyrene, a known carcinogen that has been identified in both tobacco and cannabis smoke(30, 80). The up-regulation of *MYC* combined with our finding that cannabis smoke down-regulated the tumour suppressor gene *TP53* and associated pathways suggests cannabis smoke may predispose airway epithelial cells to oncogenesis. However, it is not entirely clear why transcriptional signals associated with oncogenesis that are observed *in vitro* would not manifest into pathology *in vivo*. It is possible that the immunomodulatory effects of cannabinoids (31–34) provide protection *in vivo* via activities on additional cells beyond airway epithelial cells.

In addition to carcinogenic effects, tobacco smoke and polyaromatic hydrocarbons such as benzo-a-pyrene are known to be associated with oxidative stress via the NFE2L2/Nrf2 pathway (81–83). Similar to tobacco smoke, our transcriptome analysis of airway epithelial cells exposed to cannabis smoke indicated upregulation of the Nrf2 oxidative stress response genes *NQO1*, and *OSGIN1* in addition to a significant functional pathway enrichment of “Oxidative Stress Induced Gene Expression Via Nrf2”. We corroborated this finding by examining reactive oxygen species production in airway epithelial cells exposed to both cannabis and tobacco smokes and found that similar to tobacco smoke, cannabis smoke induced increased levels of reactive oxygen species generation, consistent with phenotypes observed in other *in vitro* systems (84). In our intervention study we found that LABA/GCS did not attenuate oxidative stress gene signatures or reactive oxygen species generation. The oxidative stress induced by cannabis smoke exposure is therefore persistent in the presence of anti-inflammatory medications used in chronic respiratory disease management, and may further impact disease development (85).

Tobacco smoke exposure experiments have used standardized research source material to ensure experimental reproducibility and robustness by limiting batch to batch variability that may impact data generation and interpretation (8, 10, 37–40). In contrast, cannabis smoke exposure experiments have not benefited from a widely accessible and chemically defined source material. For this reason, we decided to use a cannabis strain that was representative of that available in the medicinal cannabis market in Canada, that included 13% THCA (w/w), 0.18% THC, 0.35% THCVA, and 0.18% CBGA with no levels of CBD detected. Our results must therefore be interpreted based on this chemical composition and care should be taken to generalize that all cannabis strains will induce similar responses to epithelial cell barrier function, immune profile, and transcriptional activities. Indeed, increasing evidence suggests that there may be complex interactions between THC and CBD via the CB1 and CB2 cannabinoid receptors that could impact the immunomodulatory functions of cannabis smoke (31–34, 41–43). Although current law may restrict universal sharing of cannabis between research groups to ensure consistency in reagents, future cannabis smoke exposure studies could collectively benefit by reporting the chemical composition of the strain that was used to help facilitate interpretation of data generated

Our study has several limitations in experimental design that need to be highlighted. Our generation of smoke-exposure conditioned media has been based on methods described for tobacco smoke and used extensively in multiple research labs for *in vitro* studies (8, 10, 37, 38, 64). Despite this accepted method, it has not been used to create comparable smoke conditioned media extracts from different combusted materials. We therefore had to devise methods to standardize the dilution of both cannabis and tobacco smoke conditioned medias to each other to enable comparisons. Furthermore, although we normalized mass of tobacco and cannabis for cigarette composition, we opted to smoke tobacco cigarettes with filters on while cannabis cigarettes did not have a filter, as this more closely models human consumption practice. Collectively, our methods may not entirely standardize each smoke exposure dilution, representing a limitation in experiment design. Lastly, our smoke exposure experiments were limited to a single 24h exposure followed by outcome measurements. Although robust changes in epithelial cell barrier function, immune profile, and transcriptional signatures were observed in this window, the consequences of repeated cannabis smoke exposures on airway epithelial cell function should be carefully inferred from our dataset.

In conclusion, we performed a comparative study between cannabis and tobacco smoke exposure in the Calu-3 human airway epithelial cell line using concentration-response and LABA/GCS interventions with multiplex cytokine analysis and transcriptomics. Our data demonstrate striking similarities in the impacts of cannabis and tobacco smoke on airway epithelial cell barrier function, cytokine profile, and gene expression signatures. Despite the arrival of cannabis legalization, our data suggest that cannabis smoke exposure still poses a significant health risk and warrants ongoing study to build a body of clinically relevant evidence to support public policy, government regulations, and individual user practices.

## ACKNOWLEDGEMENTS

We would like to thank Dr. Jonathan Page for access to his licensed research facility and cannabis for our experiments. We would also like to thank Dr. Fiona Whelan for helping bring together the authors of the present manuscript for this collaboration. Lastly, we would like to thank Dr. James MacKillop and the McMaster University Centre for Medicinal Cannabis Research for their support of this project.

## FIGURE LEGENDS

Supplemental Figure 1. Effect of cannabis and tobacco on formoterol/budesonide-induced gene expression profile changes. Log_2_ fold change for (TSE + Form/Bud)/(Control + Form/Bud) was plotted against log_2_ fold change for (CSE + Form/Bud)/(Control + Form/Bud). High positive correlation is seen between the two smokes (Pearson correlation, r = 0.772, *p* < 1×10^−15^). 2165 genes showed significant (*q* < 0.05) expression patterns common to both cannabis and tobacco (as determined by analysis of differential expression between ((CSE + Form/Bud) + (TSE + Form/Bud)) vs (Control + Form/Bud)) and are highlighted in blue (**Supplementary Tables 10-11**). No genes were determined to be differentially expressed (*q* < 0.05) between cannabis and tobacco.

Author contributions
Jennifer A. Aguiar: Responsible for drafting the manuscript, performing the bioinformatics analysis, figure generation, and editing the manuscript.
Ryan D. Huff: Responsible for the *in vitro* experiments, *in vitro* figure generation, and editing the manuscript.
Wayne Tse: Responsible for *in vitro* experiments, and *in vitro* figure generation.
Martin R. Stampfli: Responsible for conceptual design of experiments and manuscript, identification of appropriate literature and data, and editing the manuscript.
Brendan J. McConkey: Responsible for supervision and conceptual design of bioinformatics analyses
performed.
Andrew C. Doxey: Responsible for oversight of the entire study (data collection, analysis, drafting, finalization) and supervision of the trainees
Jeremy A. Hirota: Responsible for oversight of the entire study (data collection, analysis, drafting, finalization) and supervision of the trainees

## REFERENCES

1. World Heatlh Organization - World Drug Report 2018 (United Nations publication SNEX.

2. Canada H. Canadian Cannabis Survey. 2017; http://epe.lac-bac.gc.ca/100/200/301/pwgsc-tpsgc/por-ef/health/2017/102-16-e/index.html.

3. Canada H. Canadian Cannabis Survey. 2018; http://epe.lac-bac.gc.ca/100/200/301/pwgsc-tpsgc/por-ef/health/2018/006-18-e/index.html.

4. Amatngalim GD, Schrumpf JA, Henic A, Dronkers E, Verhoosel RM, Ordonez SR, Haagsman HP, Fuentes ME, Sridhar S, Aarbiou J, Janssen RAJ, Lekkerkerker AN, Hiemstra PS. Antibacterial Defense of Human Airway Epithelial Cells from Chronic Obstructive Pulmonary Disease Patients Induced by Acute Exposure to Nontypeable Haemophilus influenzae: Modulation by Cigarette Smoke. Journal of innate immunity 2017; 9: 359–374.

5. Amatngalim GD, Broekman W, Daniel NM, van der Vlugt LE, van Schadewijk A, Taube C, Hiemstra PS. Cigarette Smoke Modulates Repair and Innate Immunity following Injury to Airway Epithelial Cells. PLoS One 2016; 11: e0166255.

6. Eddleston J, Lee RU, Doerner AM, Herschbach J, Zuraw BL. Cigarette smoke decreases innate responses of epithelial cells to rhinovirus infection. Am J Respir Cell Mol Biol 2011; 44: 118–126.

7. Thun MJ, Carter BD, Feskanich D, Freedman ND, Prentice R, Lopez AD, Hartge P, Gapstur SM. 50-year trends in smoking-related mortality in the United States. N Engl J Med 2013; 368: 351–364.

8. Rider CF, King EM, Holden NS, Giembycz MA, Newton R. Inflammatory stimuli inhibit glucocorticoid-dependent transactivation in human pulmonary epithelial cells: rescue by long-acting beta2-adrenoceptor agonists. The Journal of pharmacology and experimental therapeutics 2011; 338: 860–869.

9. Hudy MH, Traves SL, Proud D. Transcriptional and epigenetic modulation of human rhinovirus-induced CXCL10 production by cigarette smoke. Am J Respir Cell Mol Biol 2014; 50: 571–582.

10. Hudy MH, Traves SL, Wiehler S, Proud D. Cigarette smoke modulates rhinovirus-induced airway epithelial cell chemokine production. Eur Respir J 2010; 35: 1256–1263.

11. Bauer CM, Dewitte-Orr SJ, Hornby KR, Zavitz CC, Lichty BD, Stampfli MR, Mossman KL. Cigarette smoke suppresses type I interferon-mediated antiviral immunity in lung fibroblast and epithelial cells. Journal of interferon & cytokine research: the official journal of the International Society for Interferon and Cytokine Research 2008; 28: 167–179.

12. Singanayagam A, Glanville N, Girkin JL, Ching YM, Marcellini A, Porter JD, Toussaint M, Walton RP, Finney LJ, Aniscenko J, Zhu J, Trujillo-Torralbo MB, Calderazzo MA, Grainge C, Loo SL, Veerati PC, Pathinayake PS, Nichol KS, Reid AT, James PL, Solari R, Wark PAB, Knight DA, Moffatt MF, Cookson WO, Edwards MR, Mallia P, Bartlett NW, Johnston SL. Corticosteroid suppression of antiviral immunity increases bacterial loads and mucus production in COPD exacerbations. Nature communications 2018; 9: 2229.

13. Rider CF, Shah S, Miller-Larsson A, Giembycz MA, Newton R. Cytokine-Induced Loss of Glucocorticoid Function: Effect of Kinase Inhibitors, Long-Acting beta2-Adrenoceptor Agonist and Glucocorticoid Receptor Ligands. PLoS One 2015; 10: e0116773.

14. Aguiar JAT, A.; Lobb, B.; Huff, R.D.; Nguyen, J.P.; Kim, Y.; Dvorkin-Gheva, A.; Stampfli, M.R.; Doxey, A.C.; Hirota, J.A. The impact of cigarette smoke exposure, COPD, or asthma status on ABC transporter gene expression in human airway epithelial cells. Scientific reports 2019; In press.

15. Jha P, Ramasundarahettige C, Landsman V, Rostron B, Thun M, Anderson RN, McAfee T, Peto R. 21st-century hazards of smoking and benefits of cessation in the United States. N Engl J Med 2013; 368: 341–350.

16. Auerbach O, Hammond EC, Garfinkel L. Changes in bronchial epithelium in relation to cigarette smoking, 1955-1960 vs. 1970-1977. N Engl J Med 1979; 300: 381–385.

17. Spira A, Beane J, Shah V, Liu G, Schembri F, Yang X, Palma J, Brody JS. Effects of cigarette smoke on the human airway epithelial cell transcriptome. Proc Natl Acad Sci U S A 2004; 101: 10143–10148.

18. Harvey BG, Heguy A, Leopold PL, Carolan BJ, Ferris B, Crystal RG. Modification of gene expression of the small airway epithelium in response to cigarette smoking. Journal of molecular medicine 2007; 85: 39–53.

19. Tilley AE, O’Connor TP, Hackett NR, Strulovici-Barel Y, Salit J, Amoroso N, Zhou XK, Raman T, Omberg L, Clark A, Mezey J, Crystal RG. Biologic phenotyping of the human small airway epithelial response to cigarette smoking. PLoS One 2011; 6: e22798.

20. Hancox RJ, Shin HH, Gray AR, Poulton R, Sears MR. Effects of quitting cannabis on respiratory symptoms. Eur Respir J 2015; 46: 80–87.

21. Tashkin DP, Shapiro BJ, Frank IM. Acute pulmonary physiologic effects of smoked marijuana and oral (Delta)9 -tetrahydrocannabinol in healthy young men. N Engl J Med 1973; 289: 336–341.

22. Vachon L, FitzGerald MX, Solliday NH, Gould IA, Gaensler EA. Single-dose effects of marihuana smoke. Bronchial dynamics and respiratory-center sensitivity in normal subjects. N Engl J Med 1973; 288: 985–989.

23. Tashkin DP, Shapiro BJ, Frank IM. Acute effects of smoked marijuana and oral delta9-tetrahydrocannabinol on specific airway conductance in asthmatic subjects. The American review of respiratory disease 1974; 109: 420–428.

24. Tashkin DP, Shapiro BJ, Lee YE, Harper CE. Effects of smoked marijuana in experimentally induced asthma. The American review of respiratory disease 1975; 112: 377–386.

25. Wu TC, Tashkin DP, Djahed B, Rose JE. Pulmonary hazards of smoking marijuana as compared with tobacco. N Engl J Med 1988; 318: 347–351.

26. Tashkin DP, Coulson AH, Clark VA, Simmons M, Bourque LB, Duann S, Spivey GH, Gong H. Respiratory symptoms and lung function in habitual heavy smokers of marijuana alone, smokers of marijuana and tobacco, smokers of tobacco alone, and nonsmokers. The American review of respiratory disease 1987; 135: 209–216.

27. Tan WC, Lo C, Jong A, Xing L, Fitzgerald MJ, Vollmer WM, Buist SA, Sin DD, Vancouver Burden of Obstructive Lung Disease Research G. Marijuana and chronic obstructive lung disease: a population-based study. CMAJ: Canadian Medical Association journal = journal de l’Association medicale canadienne 2009; 180: 814–820.

28. Aldington S, Williams M, Nowitz M, Weatherall M, Pritchard A, McNaughton A, Robinson G, Beasley R. Effects of cannabis on pulmonary structure, function and symptoms. Thorax 2007; 62: 1058–1063.

29. Zhang LR, Morgenstern H, Greenland S, Chang SC, Lazarus P, Teare MD, Woll PJ, Orlow I, Cox B, Cannabis, Respiratory Disease Research Group of New Z, Brhane Y, Liu G, Hung RJ. Cannabis smoking and lung cancer risk: Pooled analysis in the International Lung Cancer Consortium. International journal of cancer 2015; 136: 894–903.

30. Moir D, Rickert WS, Levasseur G, Larose Y, Maertens R, White P, Desjardins S. A comparison of mainstream and sidestream marijuana and tobacco cigarette smoke produced under two machine smoking conditions. Chem Res Toxicol 2008; 21: 494–502.

31. Boggs DL, Nguyen JD, Morgenson D, Taffe MA, Ranganathan M. Clinical and Preclinical Evidence for Functional Interactions of Cannabidiol and Delta(9)-Tetrahydrocannabinol. Neuropsychopharmacology: official publication of the American College of Neuropsychopharmacology 2018; 43: 142–154.

32. Atwood BK, Mackie K. CB2: a cannabinoid receptor with an identity crisis. Br J Pharmacol 2010; 160: 467–479.

33. Pacher P, Kunos G. Modulating the endocannabinoid system in human health and disease--successes and failures. The FEBS journal 2013; 280: 1918–1943.

34. Miller AM, Stella N. CB2 receptor-mediated migration of immune cells: it can go either way. Br J Pharmacol 2008; 153: 299–308.

35. Hirota JA, Knight DA. Human airway epithelial cell innate immunity: relevance to asthma. Curr Opin Immunol 2013; 24: 740–746.

36. Parker D, Prince A. Innate immunity in the respiratory epithelium. Am J Respir Cell Mol Biol 2011; 45: 189–201.

37. Wirtz HR, Schmidt M. Acute influence of cigarette smoke on secretion of pulmonary surfactant in rat alveolar type II cells in culture. Eur Respir J 1996; 9: 24–32.

38. Huff RD, Hsu AC, Nichol KS, Jones B, Knight DA, Wark PAB, Hansbro PM, Hirota JA. Regulation of xanthine dehydrogensase gene expression and uric acid production in human airway epithelial cells. PLoS One 2017; 12: e0184260.

39. Gaschler GJ, Skrtic M, Zavitz CC, Lindahl M, Onnervik PO, Murphy TF, Sethi S, Stampfli MR. Bacteria challenge in smoke-exposed mice exacerbates inflammation and skews the inflammatory profile. Am J RespirCrit Care Med 2009; 179: 666–675.

40. Nikota JK, Shen P, Morissette MC, Fernandes K, Roos A, Chu DK, Barra NG, Iwakura Y, Kolbeck R, Humbles AA, Stampfli MR. Cigarette smoke primes the pulmonary environment to IL-1alpha/CXCR-2-dependent nontypeable Haemophilus influenzae-exacerbated neutrophilia in mice. J Immunol 2014; 193: 3134–3145.

41. Kaplan BL, Springs AE, Kaminski NE. The profile of immune modulation by cannabidiol (CBD) involves deregulation of nuclear factor of activated T cells (NFAT). Biochemical pharmacology 2008; 76: 726–737.

42. Mecha M, Feliu A, Inigo PM, Mestre L, Carrillo-Salinas FJ, Guaza C. Cannabidiol provides long-lasting protection against the deleterious effects of inflammation in a viral model of multiple sclerosis: a role for A2A receptors. Neurobiology of disease 2013; 59: 141–150.

43. Srivastava MD, Srivastava BI, Brouhard B. Delta9 tetrahydrocannabinol and cannabidiol alter cytokine production by human immune cells. Immunopharmacology 1998; 40: 179–185.

44. Barnes NC, Qiu YS, Pavord ID, Parker D, Davis PA, Zhu J, Johnson M, Thomson NC, Jeffery PK, Group SCOS. Antiinflammatory effects of salmeterol/fluticasone propionate in chronic obstructive lung disease. Am J Respir Crit Care Med 2006; 173: 736–743.

45. Kew KM, Dias S, Cates CJ. Long-acting inhaled therapy (beta-agonists, anticholinergics and steroids) for COPD: a network meta-analysis. The Cochrane database of systematic reviews 2014: CD010844.

46. Calverley P, Pauwels R, Vestbo J, Jones P, Pride N, Gulsvik A, Anderson J, Maden C, STeroids TRoI, long-acting beta2 agonists study g. Combined salmeterol and fluticasone in the treatment of chronic obstructive pulmonary disease: a randomised controlled trial. Lancet 2003; 361: 449–456.

47. Pauwels RA, Lofdahl CG, Postma DS, Tattersfield AE, O’Byrne P, Barnes PJ, Ullman A. Effect of inhaled formoterol and budesonide on exacerbations of asthma. Formoterol and Corticosteroids Establishing Therapy (FACET) International Study Group. N Engl J Med 1997; 337: 1405–1411.

48. Giembycz MA, Kaur M, Leigh R, Newton R. A Holy Grail of asthma management: toward understanding how long-acting beta(2)-adrenoceptor agonists enhance the clinical efficacy of inhaled corticosteroids. Br J Pharmacol 2008; 153: 1090–1104.

49. Giembycz MA, Newton R. Potential mechanisms to explain how LABAs and PDE4 inhibitors enhance the clinical efficacy of glucocorticoids in inflammatory lung diseases. F1000prime reports 2015; 7: 16.

50. Kaur M, Chivers JE, Giembycz MA, Newton R. Long-acting beta2-adrenoceptor agonists synergistically enhance glucocorticoid-dependent transcription in human airway epithelial and smooth muscle cells. Molecular pharmacology 2008; 73: 203–214.

51. Kreft ME, Jerman UD, Lasic E, Hevir-Kene N, Rizner TL, Peternel L, Kristan K. The characterization of the human cell line Calu-3 under different culture conditions and its use as an optimized in vitro model to investigate bronchial epithelial function. European journal of pharmaceutical sciences: official journal of the European Federation for Pharmaceutical Sciences 2015; 69: 1–9.

52. Huff RD, Rider CF, Yan D, Newton R, Giembycz MA, Carlsten C, Hirota JA. Inhibition of ABCC4 potentiates combination beta agonist and glucocorticoid responses in human airway epithelial cells. J Allergy Clin Immunol 2017.

53. Schlafli AM, Adams O, Galvan JA, Gugger M, Savic S, Bubendorf L, Schmid RA, Becker KF, Tschan MP, Langer R, Berezowska S. Prognostic value of the autophagy markers LC3 and p62/SQSTM1 in early-stage non-small cell lung cancer. Oncotarget 2016; 7: 39544–39555.

54. Hussain T, Al-Attas OS, Al-Daghri NM, Mohammed AA, De Rosas E, Ibrahim S, Vinodson B, Ansari MG, El-Din KI. Induction of CYP1A1, CYP1A2, CYP1B1, increased oxidative stress and inflammation in the lung and liver tissues of rats exposed to incense smoke. Molecular and cellular biochemistry 2014; 391: 127–136.

55. Zuo J, Brewer DS, Arlt VM, Cooper CS, Phillips DH. Benzo pyrene-induced DNA adducts and gene expression profiles in target and non-target organs for carcinogenesis in mice. BMC Genomics 2014; 15: 880.

56. van Delft J, Gaj S, Lienhard M, Albrecht MW, Kirpiy A, Brauers K, Claessen S, Lizarraga D, Lehrach H, Herwig R, Kleinjans J. RNA-Seq provides new insights in the transcriptome responses induced by the carcinogen benzo[a]pyrene. Toxicol Sci 2012; 130: 427–439.

57. van Delft JH, Mathijs K, Staal YC, van Herwijnen MH, Brauers KJ, Boorsma A, Kleinjans JC. Time series analysis of benzo[A]pyrene-induced transcriptome changes suggests that a network of transcription factors regulates the effects on functional gene sets. Toxicol Sci 2010; 117: 381–392.

58. Barouki R, Morel Y. Repression of cytochrome P450 1A1 gene expression by oxidative stress: mechanisms and biological implications. Biochemical pharmacology 2001; 61: 511–516.

59. Cho HY, Reddy SP, Kleeberger SR. Nrf2 defends the lung from oxidative stress. Antioxid Redox Signal 2006; 8: 76–87.

60. Luzina IG, Lockatell V, Hyun SW, Kopach P, Kang PH, Noor Z, Liu A, Lillehoj EP, Lee C, Miranda-Ribera A, Todd NW, Goldblum SE, Atamas SP. Elevated expression of NEU1 sialidase in idiopathic pulmonary fibrosis provokes pulmonary collagen deposition, lymphocytosis, and fibrosis. Am J Physiol Lung Cell Mol Physiol 2016; 310: L940–954.

61. Kicic A, Sutanto EN, Stevens PT, Knight DA, Stick SM. Intrinsic biochemical and functional differences in bronchial epithelial cells of children with asthma. Am J RespirCrit Care Med 2006; 174: 1110–1118.

62. Steiling K, van den Berge M, Hijazi K, Florido R, Campbell J, Liu G, Xiao J, Zhang X, Duclos G, Drizik E, Si H, Perdomo C, Dumont C, Coxson HO, Alekseyev YO, Sin D, Pare P, Hogg JC, McWilliams A, Hiemstra PS, Sterk PJ, Timens W, Chang JT, Sebastiani P, O’Connor GT, Bild AH, Postma DS, Lam S, Spira A, Lenburg ME. A dynamic bronchial airway gene expression signature of chronic obstructive pulmonary disease and lung function impairment. Am J Respir Crit Care Med 2013; 187: 933–942.

63. Xu Y, Mizuno T, Sridharan A, Du Y, Guo M, Tang J, Wikenheiser-Brokamp KA, Perl AT, Funari VA, Gokey JJ, Stripp BR, Whitsett JA. Single-cell RNA sequencing identifies diverse roles of epithelial cells in idiopathic pulmonary fibrosis. JCI insight 2016; 1: e90558.

64. Schamberger AC, Mise N, Jia J, Genoyer E, Yildirim AO, Meiners S, Eickelberg O. Cigarette smoke-induced disruption of bronchial epithelial tight junctions is prevented by transforming growth factor-beta. Am J Respir Cell Mol Biol 2014; 50: 1040–1052.

65. Moheimani F, Roth HM, Cross J, Reid AT, Shaheen F, Warner SM, Hirota JA, Kicic A, Hallstrand TS, Kahn M, Stick SM, Hansbro PM, Hackett TL, Knight DA. Disruption of beta-catenin/CBP signaling inhibits human airway epithelial-mesenchymal transition and repair. Int J Biochem Cell Biol 2015; 68: 59–69.

66. Emami KH, Nguyen C, Ma H, Kim DH, Jeong KW, Eguchi M, Moon RT, Teo JL, Kim HY, Moon SH, Ha JR, Kahn M. A small molecule inhibitor of beta-catenin/CREB-binding protein transcription [corrected]. Proc Natl Acad Sci U S A 2004; 101: 12682–12687.

67. Goubau D, Deddouche S, Reis e Sousa C. Cytosolic sensing of viruses. Immunity 2013; 38: 855–869.

68. Modestou MA, Manzel LJ, El-Mahdy S, Look DC. Inhibition of IFN-gamma-dependent antiviral airway epithelial defense by cigarette smoke. Respir Res 2010; 11: 64.

69. Pawliczak R, Logun C, Madara P, Barb J, Suffredini AF, Munson PJ, Danner RL, Shelhamer JH. Influence of IFN-gamma on gene expression in normal human bronchial epithelial cells: modulation of IFN-gamma effects by dexamethasone. Physiol Genomics 2005; 23: 28–45.

70. Hogg JC, Pare PD, Hackett TL. The Contribution of Small Airway Obstruction to the Pathogenesis of Chronic Obstructive Pulmonary Disease. Physiological reviews 2017; 97: 529–552.

71. Janson C, Stratelis G, Miller-Larsson A, Harrison TW, Larsson K. Scientific rationale for the possible inhaled corticosteroid intraclass difference in the risk of pneumonia in COPD. International journal of chronic obstructive pulmonary disease 2017; 12: 3055–3064.

72. Drummond MB, Dasenbrook EC, Pitz MW, Murphy DJ, Fan E. Inhaled corticosteroids in patients with stable chronic obstructive pulmonary disease: a systematic review and meta-analysis. Jama 2008; 300: 2407–2416.

73. Morjaria JB, Rigby A, Morice AH. Inhaled Corticosteroid use and the Risk of Pneumonia and COPD Exacerbations in the UPLIFT Study. Lung 2017; 195: 281–288.

74. Skevaki CL, Christodoulou I, Spyridaki IS, Tiniakou I, Georgiou V, Xepapadaki P, Kafetzis DA, Papadopoulos NG. Budesonide and formoterol inhibit inflammatory mediator production by bronchial epithelial cells infected with rhinovirus. Clin Exp Allergy 2009; 39: 1700–1710.

75. Kurt OK, Zhang J, Pinkerton KE. Pulmonary health effects of air pollution. Current opinion in pulmonary medicine 2016; 22: 138–143.

76. Gordon SB, Bruce NG, Grigg J, Hibberd PL, Kurmi OP, Lam KB, Mortimer K, Asante KP, Balakrishnan K, Balmes J, Bar-Zeev N, Bates MN, Breysse PN, Buist S, Chen Z, Havens D, Jack D, Jindal S, Kan H, Mehta S, Moschovis P, Naeher L, Patel A, Perez-Padilla R, Pope D, Rylance J, Semple S, Martin WJ, 2nd. Respiratory risks from household air pollution in low and middle income countries. The Lancet Respiratory medicine 2014; 2: 823–860.

77. Thakur M, Nuyts PAW, Boudewijns EA, Flores Kim J, Faber T, Babu GR, van Schayck OCP, Been JV. Impact of improved cookstoves on women’s and child health in low and middle income countries: a systematic review and meta-analysis. Thorax 2018; 73: 1026–1040.

78. Kc R, Shukla SD, Gautam SS, Hansbro PM, O’Toole RF. The role of environmental exposure to non-cigarette smoke in lung disease. Clinical and translational medicine 2018; 7: 39.

79. Lu L, Qi H, Luo F, Xu H, Ling M, Qin Y, Yang P, Liu X, Yang Q, Xue J, Chen C, Lu J, Xiang Q, Liu Q, Bian Q. Feedback circuitry via let-7c between lncRNA CCAT1 and c-Myc is involved in cigarette smoke extract-induced malignant transformation of HBE cells. Oncotarget 2017; 8: 19285–19297.

80. Fields WR, Desiderio JG, Leonard RM, Burger EE, Brown BG, Doolittle DJ. Differential c-myc expression profiles in normal human bronchial epithelial cells following treatment with benzo[a]pyrene, benzo[a]pyrene-4,5 epoxide, and benzo[a]pyrene-7,8-9,10 diol epoxide. Molecular carcinogenesis 2004; 40: 79–89.

81. Valdivieso AG, Dugour AV, Sotomayor V, Clauzure M, Figueroa JM, Santa-Coloma TA. N-acetyl cysteine reverts the proinflammatory state induced by cigarette smoke extract in lung Calu-3 cells. Redox biology 2018; 16: 294–302.

82. Faux SP, Tai T, Thorne D, Xu Y, Breheny D, Gaca M. The role of oxidative stress in the biological responses of lung epithelial cells to cigarette smoke. Biomarkers: biochemical indicators of exposure, response, and susceptibility to chemicals 2009; 14 Suppl 1: 90–96.

83. Nguyen PM, Park MS, Chow M, Chang JH, Wrischnik L, Chan WK. Benzo[a]pyrene increases the Nrf2 content by downregulating the Keap1 message. Toxicol Sci 2010; 116: 549–561.

84. Sarafian TA, Magallanes JA, Shau H, Tashkin D, Roth MD. Oxidative stress produced by marijuana smoke. An adverse effect enhanced by cannabinoids. Am J Respir Cell Mol Biol 1999; 20: 1286–1293.

85. Park HS, Kim SR, Lee YC. Impact of oxidative stress on lung diseases. Respirology 2009; 14: 27–38.

