## Supplement Figure 1 for "A comparative analysis of cannabis and tobacco smoke exposure on human airway epithelial cell gene expression, immune phenotype, and response to formoterol and budesonide treatment"

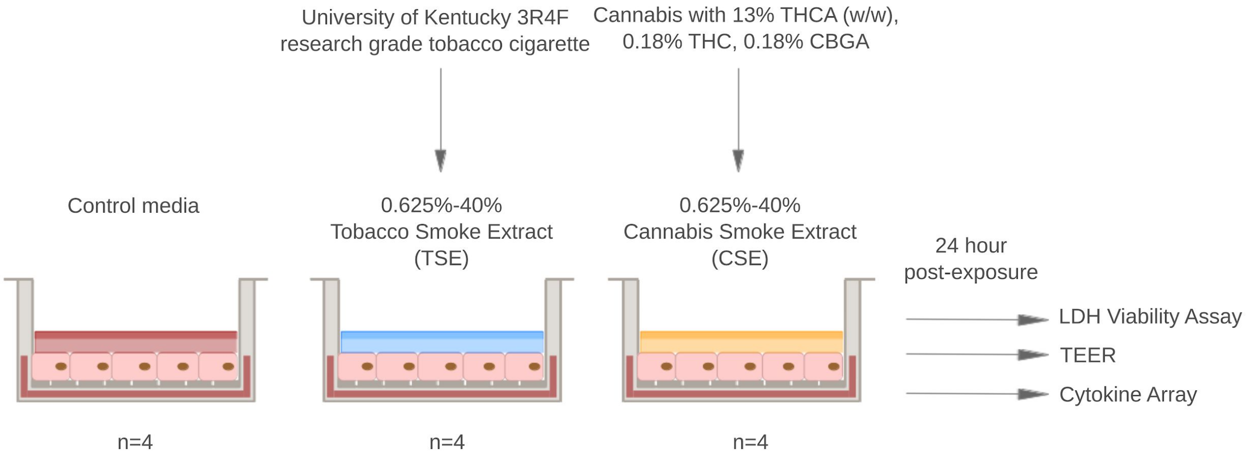
University of Kentucky 3R4F
research grade tobacco cigarette
Cannabis with 13% THCA (w/w),
0.18% THC, 0.18% CBGA
Control media
0.625%-40%
Tobacco Smoke Extract
(TSE)
0.625%-40%
Cannabis Smoke Extract
(CSE)
24 hour
post-exposure
LDH Viability Assay
TEER
Cytokine Array
n=4
n=4
n=4

Figure 2

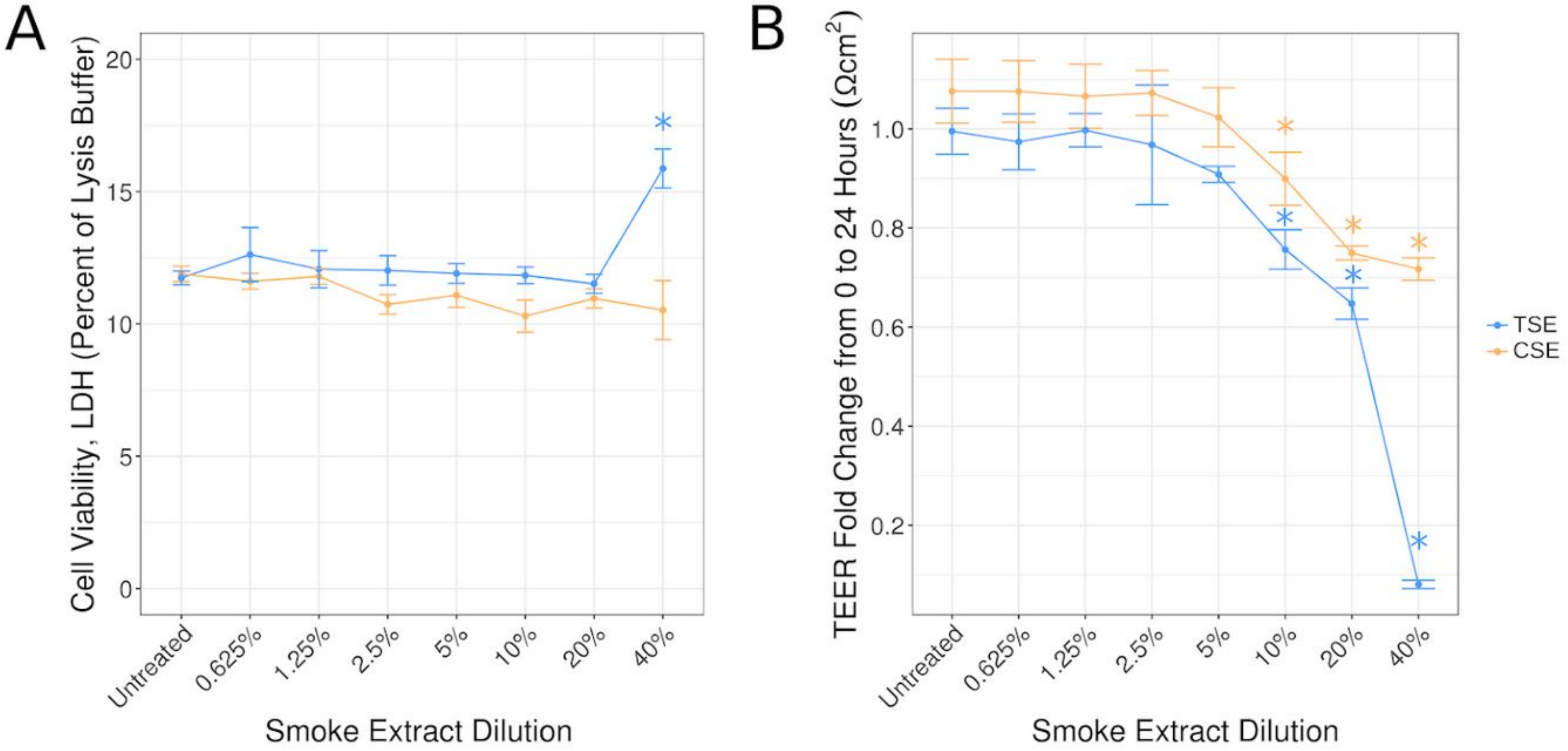
A
Cell Viability, LDH (Percent of Lysis Buffer)
20
15
10
5
0
*
Untreated
0.625%
1.25%
2.5%
5%
10%
20%
40%
Smoke Extract Dilution
B
TEER Fold Change from 0 to 24 Hours (Ωcm²)
1.0
0.8
0.6
0.4
0.2
Untreated
0.625%
1.25%
2.5%
5%
10%
20%
40%
Smoke Extract Dilution
TSE
CSE

Figure 3

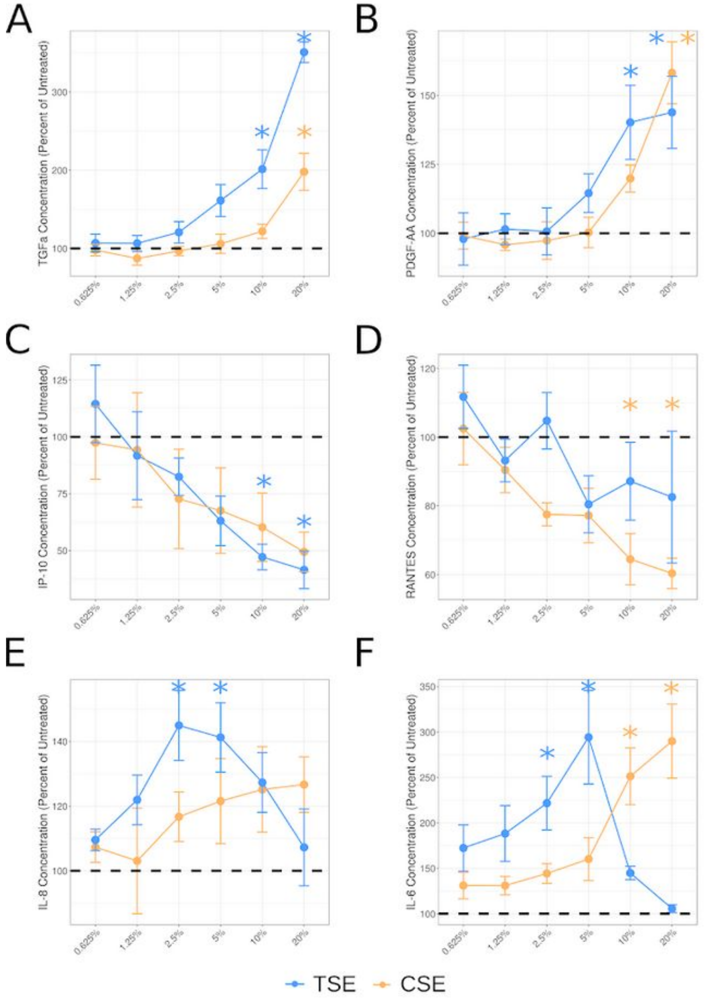
A
B
C
D
E
F
TGFa Concentration (Percent of Untreated)
PDGF-AA Concentration (Percent of Untreated)
IP-10 Concentration (Percent of Untreated)
RANTES Concentration (Percent of Untreated)
IL-8 Concentration (Percent of Untreated)
IL-6 Concentration (Percent of Untreated)
0.625%
1.25%
2.5%
5%
10%
20%
TSE
CSE

### Figure 4

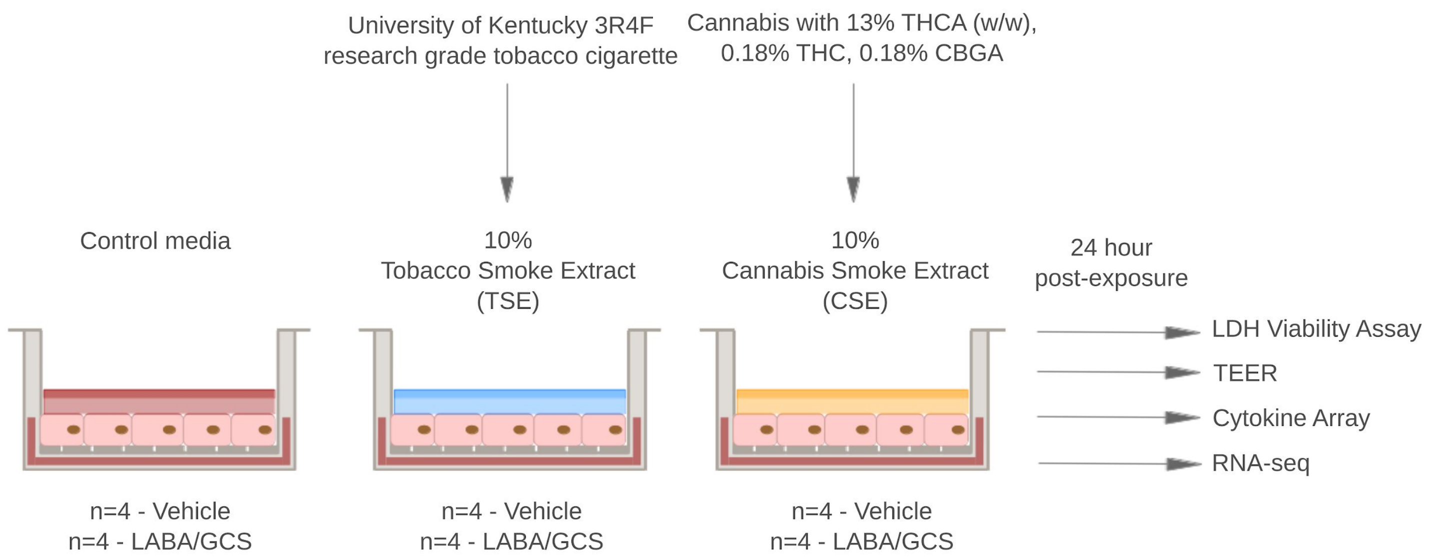

University of Kentucky 3R4F
research grade tobacco cigarette
Cannabis with 13% THCA (w/w),
0.18% THC, 0.18% CBGA
Control media
10%
Tobacco Smoke Extract
(TSE)
10%
Cannabis Smoke Extract
(CSE)
24 hour
post-exposure
LDH Viability Assay
TEER
Cytokine Array
RNA-seq
n=4 - Vehicle
n=4 - LABA/GCS
n=4 - Vehicle
n=4 - LABA/GCS
n=4 - Vehicle
n=4 - LABA/GCS

Figure 5

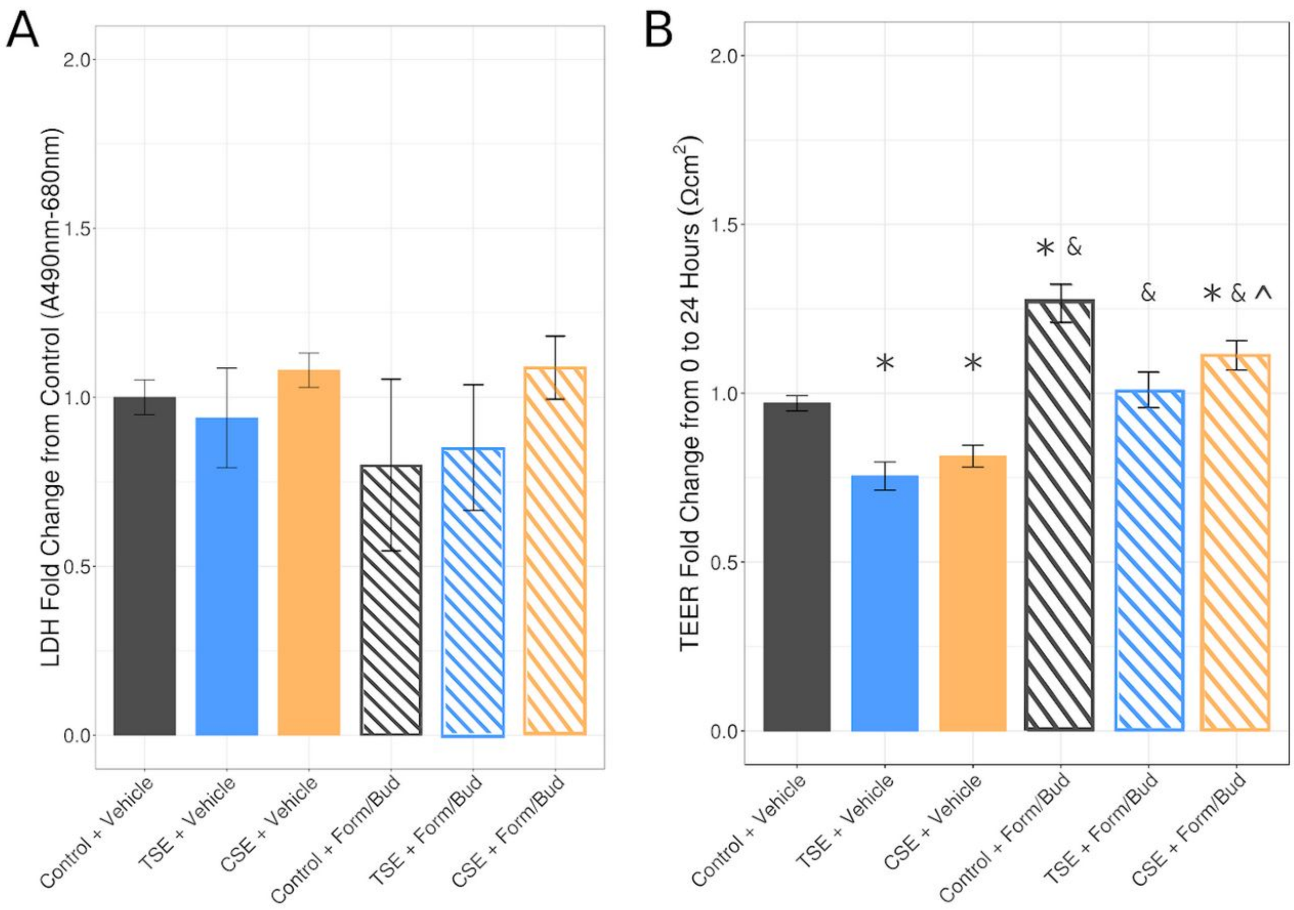

A
LDH Fold Change from Control (A490nm-680nm)
2.0
1.5
1.0
0.5
0.0
Control + Vehicle
TSE + Vehicle
CSE + Vehicle
Control + Form/Bud
TSE + Form/Bud
CSE + Form/Bud
B
TEER Fold Change from 0 to 24 Hours (Ωcm²)
2.0
1.5
1.0
0.5
0.0
*
*
* &
&
* & ^
Control + Vehicle
TSE + Vehicle
CSE + Vehicle
Control + Form/Bud
TSE + Form/Bud
CSE + Form/Bud

Figure 6

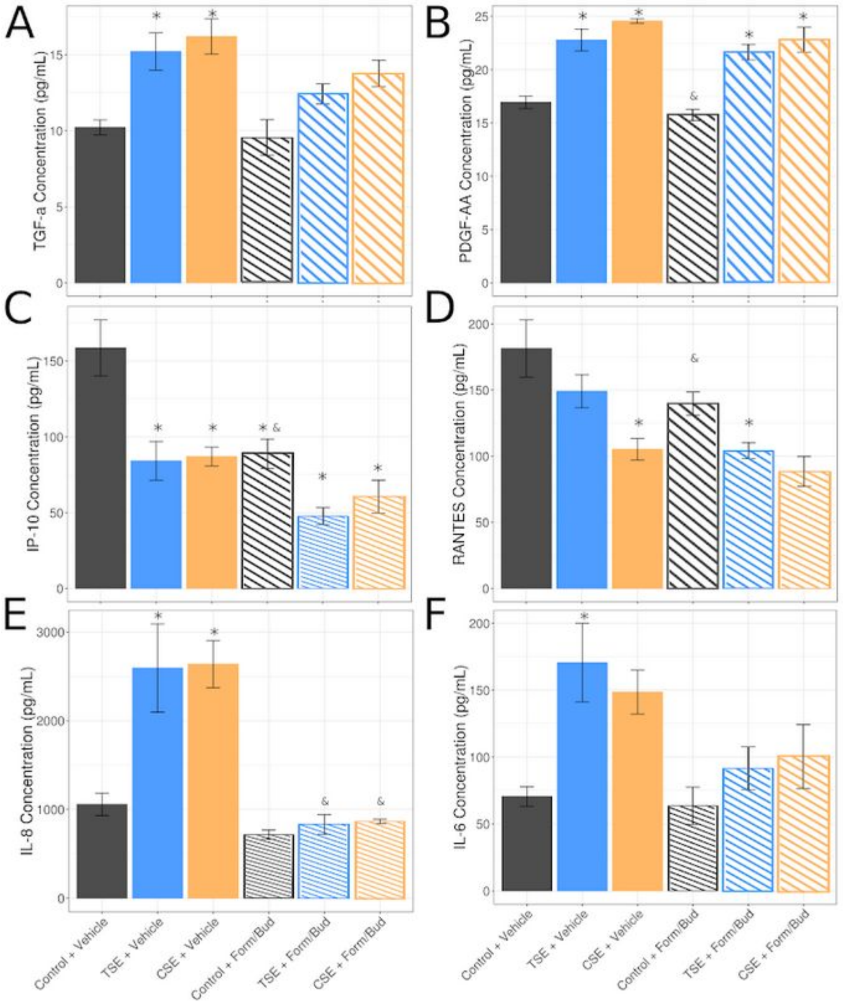
A
B
C
D
E
F
TGF-a Concentration (pg/mL)
PDGF-AA Concentration (pg/mL)
IP-10 Concentration (pg/mL)
RANTES Concentration (pg/mL)
IL-8 Concentration (pg/mL)
IL-6 Concentration (pg/mL)
Control + Vehicle
TSE + Vehicle
CSE + Vehicle
Control + Form/Bud
TSE + Form/Bud
CSE + Form/Bud

Figure 7

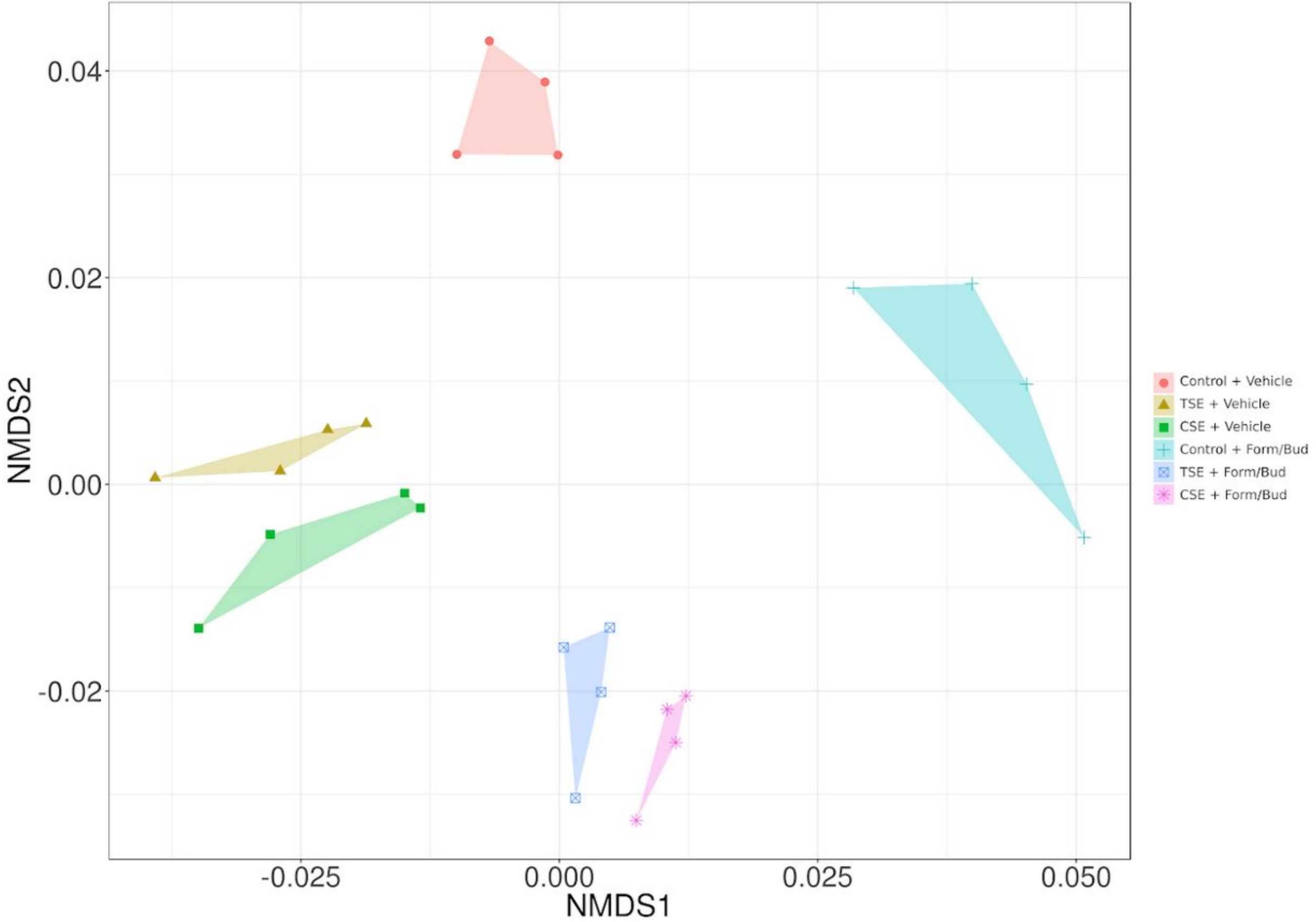
0.04
0.02
0.00
-0.02
NMDS2
-0.025
0.000
0.025
0.050
NMDS1
Control + Vehicle
TSE + Vehicle
CSE + Vehicle
Control + Form/Bud
TSE + Form/Bud
CSE + Form/Bud

Figure 8

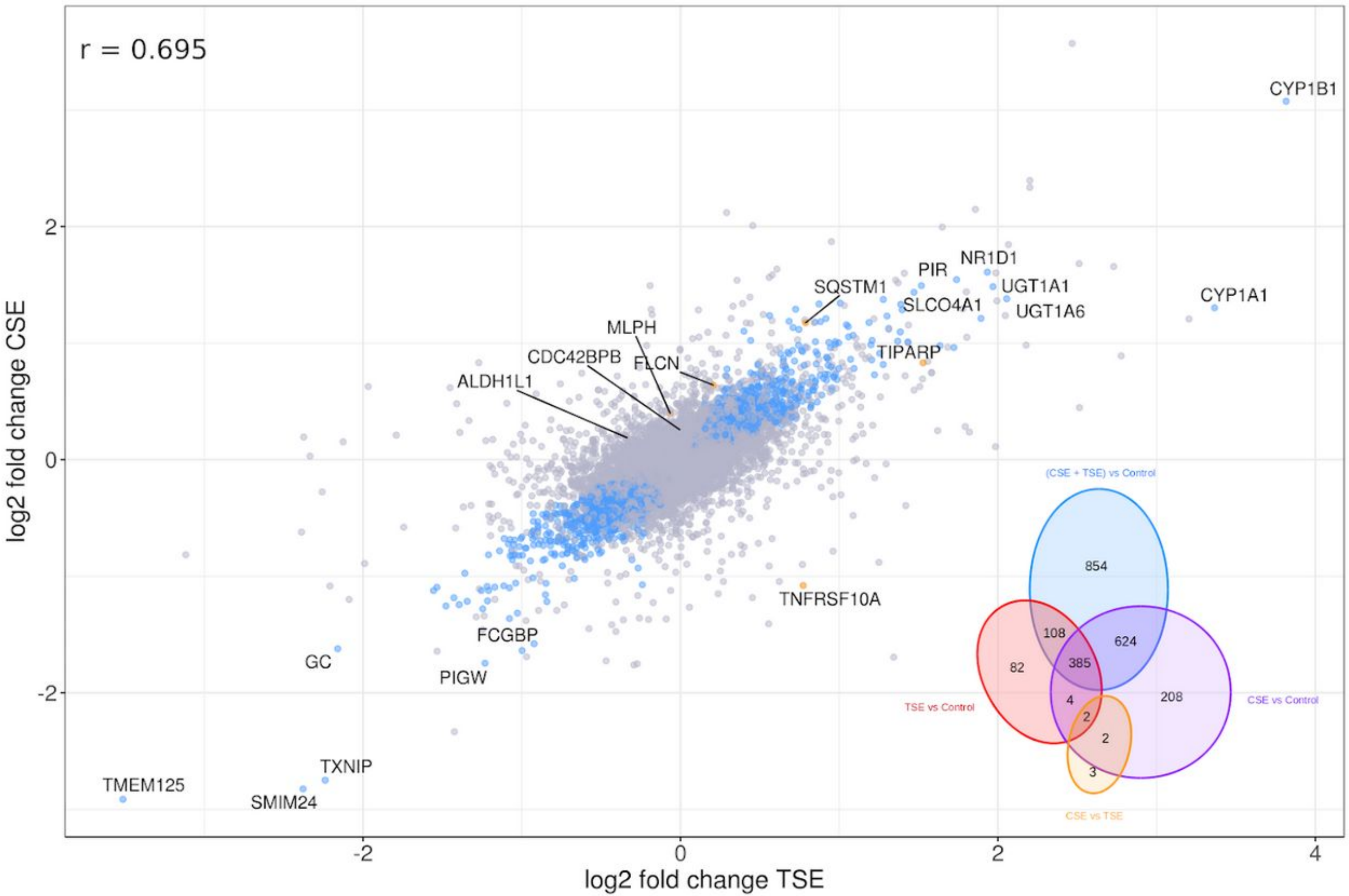

r = 0.695
CYP1B1
CYP1A1
NR1D1
PIR
UGT1A1
UGT1A6
SQSTM1
SLCO4A1
TIPARP
MLPH
CDC42BPB
FLCN
ALDH1L1
TNFRSF10A
FCGBP
GC
PIGW
TXNIP
TMEM125
SMIM24
(CSE + TSE) vs Control
854
108
624
82
385
208
4
2
2
3
TSE vs Control
CSE vs Control
CSE vs TSE
2
0
-2
-2
0
2
4
log2 fold change CSE
log2 fold change TSE

Figure X (Omitted in BioRxiv Preprint)

- Define network based on B[a]p signature genes
- Color nodes by log2 fold change
- Point of figure can be that B[a]p induced genes form an interconnected network and that the network responds almost identically between both smoke exposures, implicating B[a]p as the main driver of the response

A Cannabis

B Tobacco

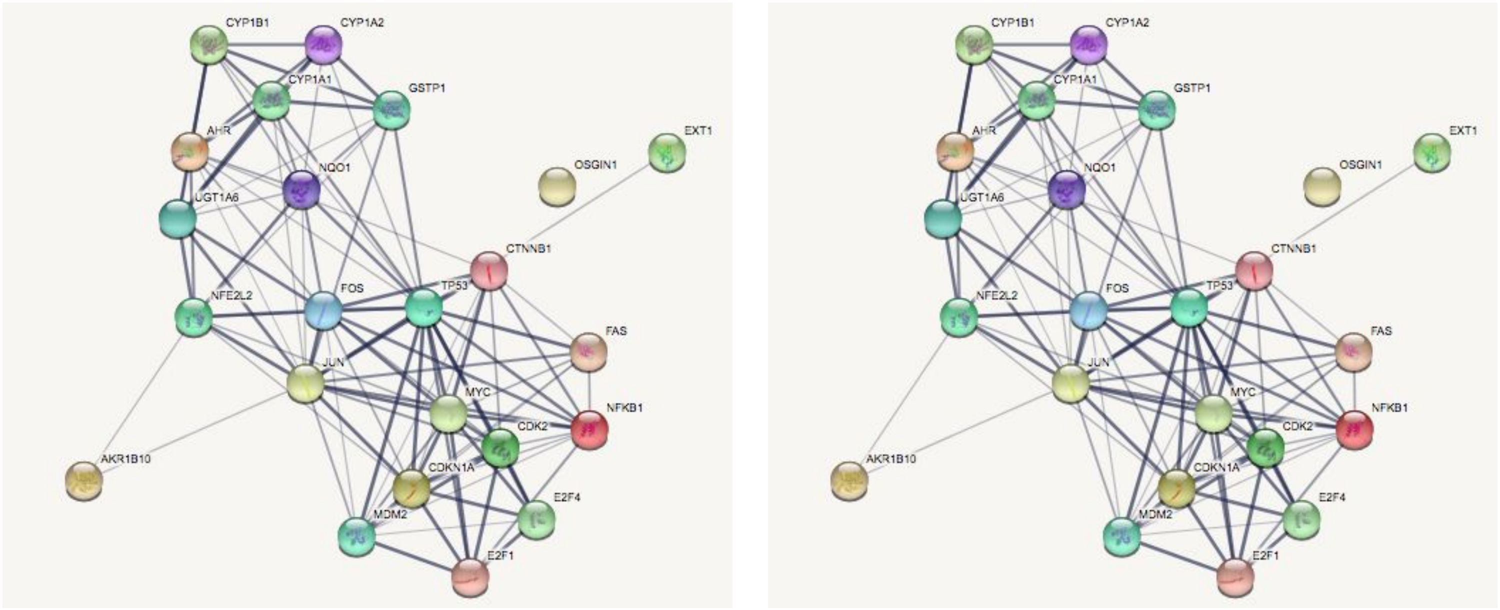

Figure 9

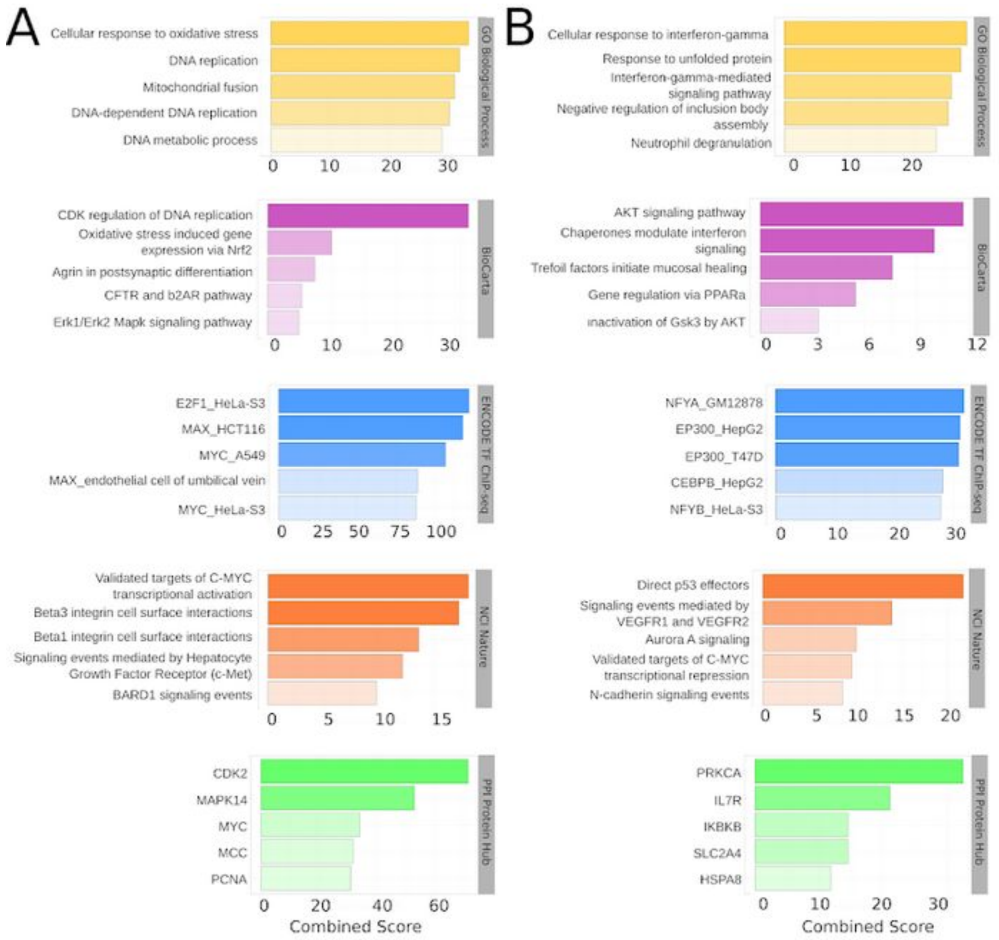
A
Cellular response to oxidative stress
DNA replication
Mitochondrial fusion
DNA-dependent DNA replication
DNA metabolic process
GO Biological Process
CDK regulation of DNA replication
Oxidative stress induced gene expression via Nrf2
Agrin in postsynaptic differentiation
CFTR and b2AR pathway
Erk1/Erk2 Mapk signaling pathway
BioCarta
E2F1_HeLa-S3
MAX_HCT116
MYC_A549
MAX_endothelial cell of umbilical vein
MYC_HeLa-S3
ENCODE TF ChIP-seq
Validated targets of C-MYC transcriptional activation
Beta3 integrin cell surface interactions
Beta1 integrin cell surface interactions
Signaling events mediated by Hepatocyte Growth Factor Receptor (c-Met)
BARD1 signaling events
NCI Nature
CDK2
MAPK14
MYC
MCC
PCNA
PPI Protein Hub
Combined Score
B
Cellular response to interferon-gamma
Response to unfolded protein
Interferon-gamma-mediated signaling pathway
Negative regulation of inclusion body assembly
Neutrophil degranulation
GO Biological Process
AKT signaling pathway
Chaperones modulate interferon signaling
Trefoil factors initiate mucosal healing
Gene regulation via PPARa
inactivation of Gsk3 by AKT
BioCarta
NFYA_GM12878
EP300_HepG2
EP300_T47D
CEBPB_HepG2
NFYB_HeLa-S3
ENCODE TF ChIP-seq
Direct p53 effectors
Signaling events mediated by VEGFR1 and VEGFR2
Aurora A signaling
Validated targets of C-MYC transcriptional repression
N-cadherin signaling events
NCI Nature
PRKCA
IL7R
IKBKB
SLC2A4
HSPA8
PPI Protein Hub
Combined Score

Figure 10

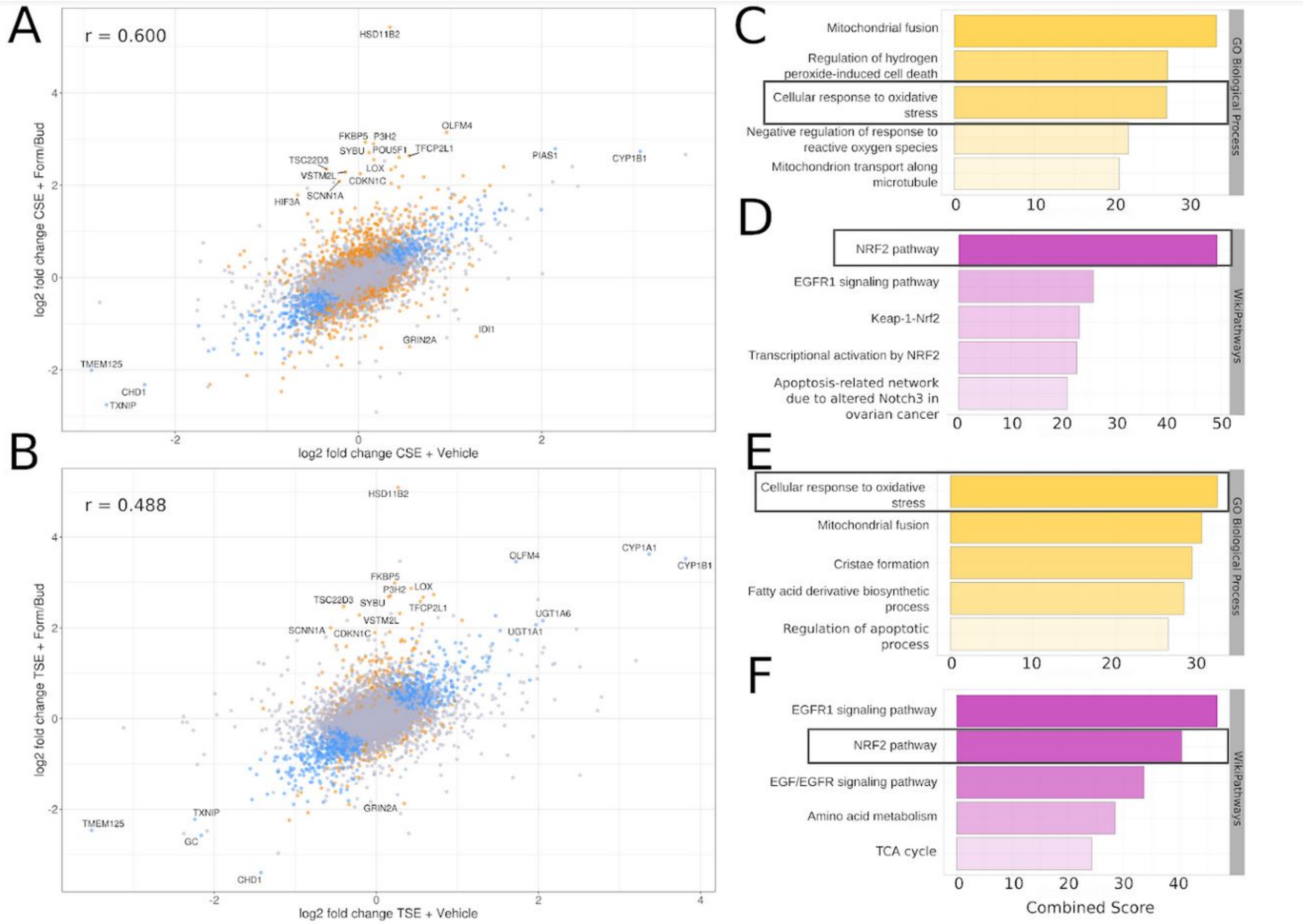
A
r = 0.600
HSD11B2
OLFM4
FKBP5
P3H2
SYBU
POU5F1
TFCP2L1
TSC22D3
VSTM2L
LOX
CDKN1C
SCNN1A
HIF3A
PIAS1
CYP1B1
IDI1
GRIN2A
TMEM125
CHD1
TXNIP
log2 fold change CSE + Form/Bud
log2 fold change CSE + Vehicle
B
r = 0.488
HSD11B2
CYP1A1
CYP1B1
OLFM4
FKBP5
P3H2
LOX
TSC22D3
SYBU
TFCP2L1
VSTM2L
SCNN1A
CDKN1C
UGT1A6
UGT1A1
GRIN2A
TXNIP
TMEM125
GC
CHD1
log2 fold change TSE + Form/Bud
log2 fold change TSE + Vehicle
C
Mitochondrial fusion
Regulation of hydrogen peroxide-induced cell death
Cellular response to oxidative stress
Negative regulation of response to reactive oxygen species
Mitochondrion transport along microtubule
GO Biological Process
D
NRF2 pathway
EGFR1 signaling pathway
Keap-1-Nrf2
Transcriptional activation by NRF2
Apoptosis-related network due to altered Notch3 in ovarian cancer
WikiPathways
E
Cellular response to oxidative stress
Mitochondrial fusion
Cristae formation
Fatty acid derivative biosynthetic process
Regulation of apoptotic process
GO Biological Process
F
EGFR1 signaling pathway
NRF2 pathway
EGF/EGFR signaling pathway
Amino acid metabolism
TCA cycle
WikiPathways
Combined Score

Figure 11

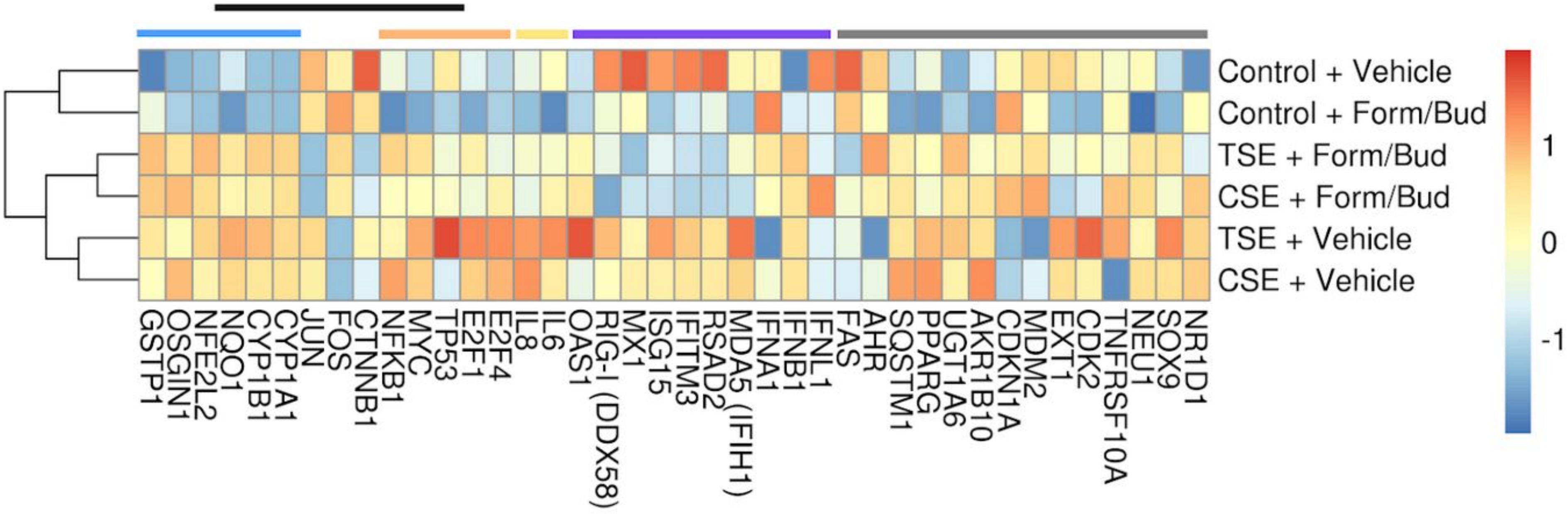

Control + Vehicle
Control + Form/Bud
TSE + Form/Bud
CSE + Form/Bud
TSE + Vehicle
CSE + Vehicle
1
0
-1
GSTP1
OSGIN1
NFE2L2
NQO1
CYP1B1
CYP1A1
JUN
FOS
CTNNB1
NFKB1
MYC
TP53
E2F1
E2F4
IL8
IL6
OAS1
RIG-I (DDX58)
MX1
ISG15
IFITM3
RSAD2
MDA5 (IFIH1)
IFNA1
IFNB1
IFNL1
FAS
AHR
SQSTM1
PPARG
UGT1A6
AKR1B10
CDKN1A
MDM2
EXT1
CDK2
TNFRSF10A
NEU1
SOX9
NR1D1

Figure 12

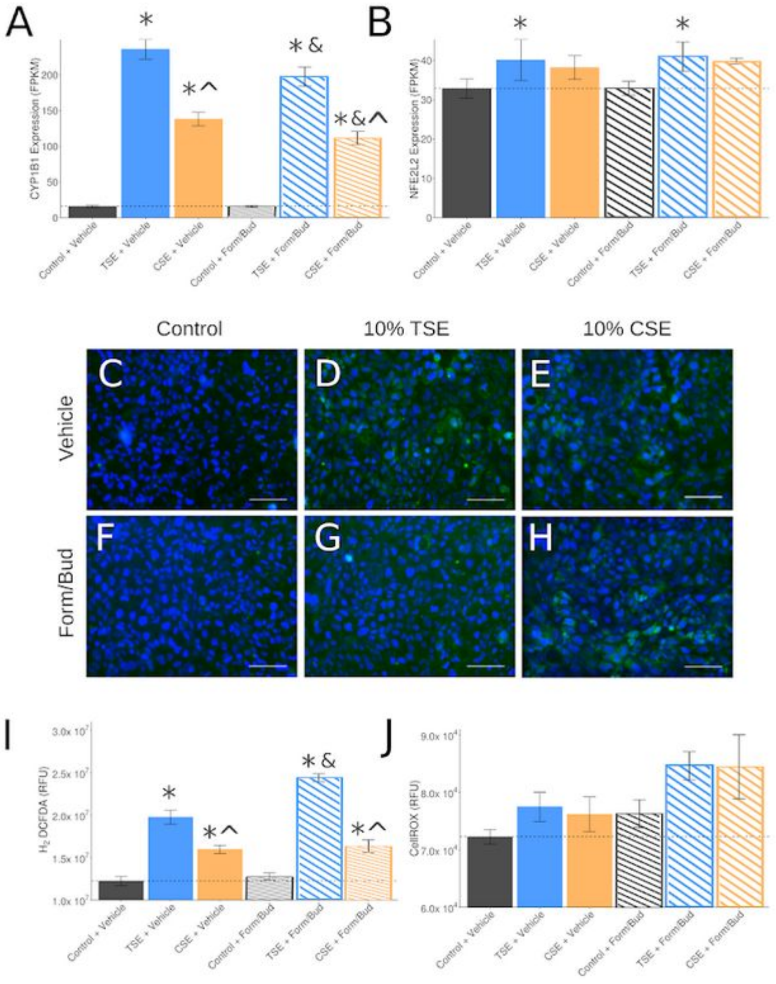

A
B
CYP1B1 Expression (FPKM)
NFE2L2 Expression (FPKM)
Control + Vehicle
TSE + Vehicle
CSE + Vehicle
Control + Form/Bud
TSE + Form/Bud
CSE + Form/Bud
Control
10% TSE
10% CSE
Vehicle
Form/Bud
C
D
E
F
G
H
I
J
H2 DCFDA (RFU)
CellROX (RFU)

Figure X (Omitted from BioRxiv Preprint)

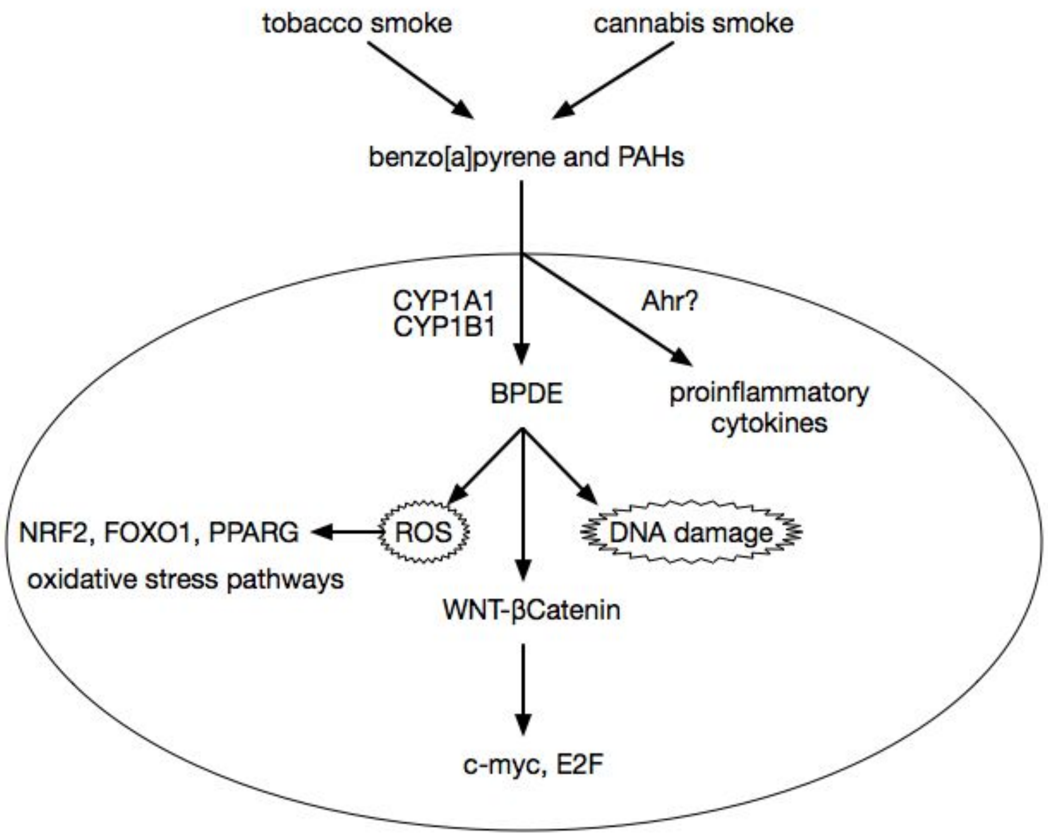

Supplement Figure 1

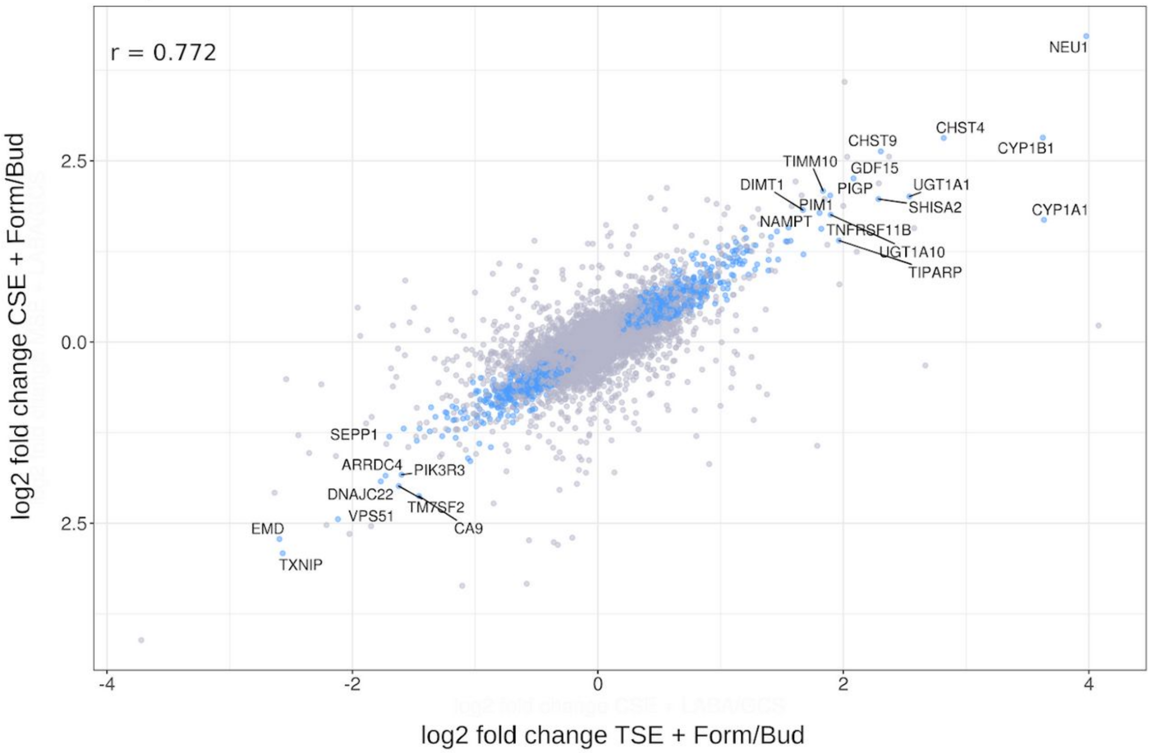

### EXTRA SLIDES/ NOTES

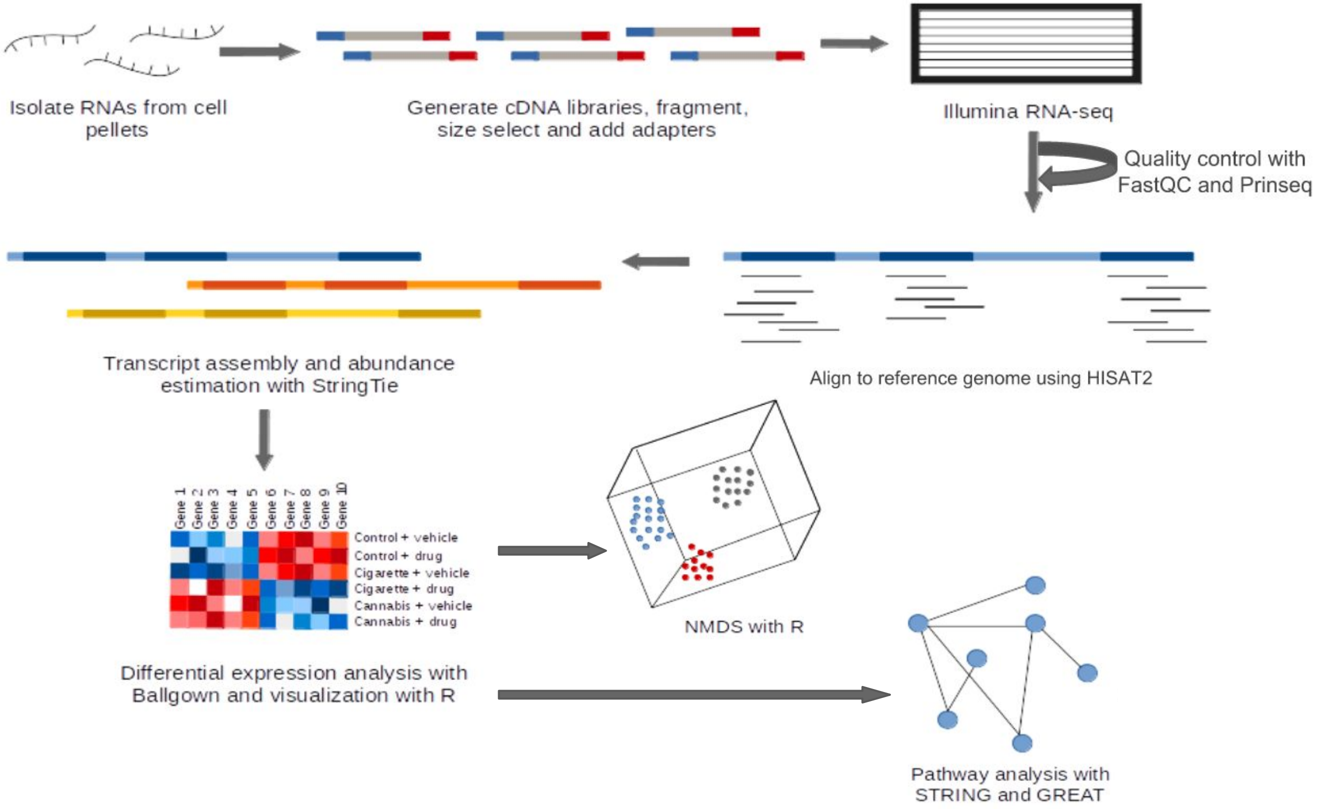

Isolate RNAs from cell pellets
Generate cDNA libraries, fragment, size select and add adapters
Illumina RNA-seq
Quality control with FastQC and Prinseq
Align to reference genome using HISAT2
Transcript assembly and abundance estimation with StringTie
Gene 1
Gene 2
Gene 3
Gene 4
Gene 5
Gene 6
Gene 7
Gene 8
Gene 9
Gene 10
Control + vehicle
Control + drug
Cigarette + vehicle
Cigarette + drug
Cannabis + vehicle
Cannabis + drug
Differential expression analysis with Ballgown and visualization with R
NMDS with R
Pathway analysis with STRING and GREAT

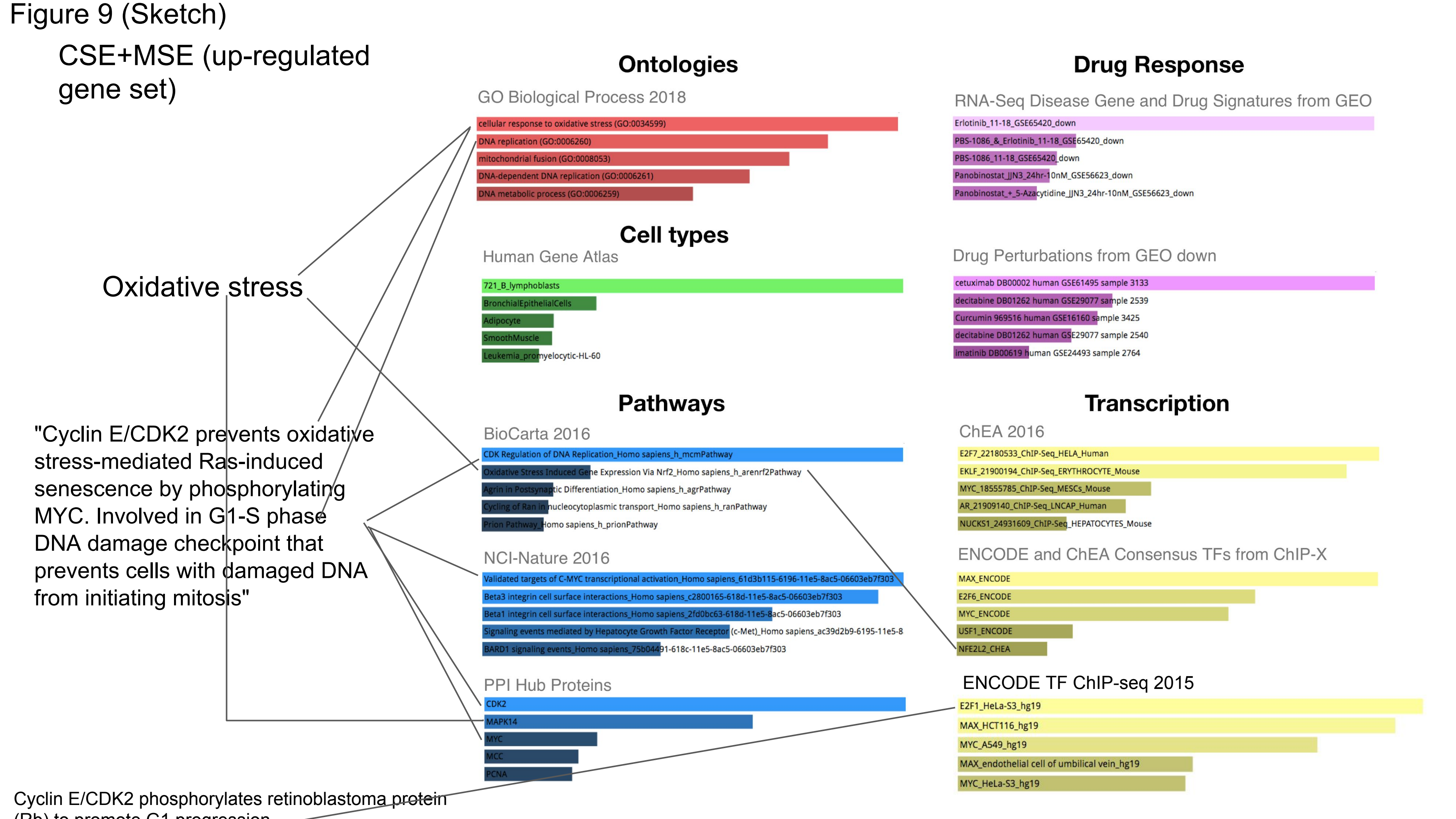

Figure 9 (Sketch)
CSE+MSE (up-regulated gene set)
Oxidative stress
"Cyclin E/CDK2 prevents oxidative stress-mediated Ras-induced senescence by phosphorylating MYC. Involved in G1-S phase DNA damage checkpoint that prevents cells with damaged DNA from initiating mitosis"
Cyclin E/CDK2 phosphorylates retinoblastoma protein (Rb) to promote G1 progression
Ontologies
GO Biological Process 2018
cellular response to oxidative stress (GO:0034599)
DNA replication (GO:0006260)
mitochondrial fusion (GO:0008053)
DNA-dependent DNA replication (GO:0006261)
DNA metabolic process (GO:0006259)
Cell types
Human Gene Atlas
721_B_lymphoblasts
BronchialEpithelialCells
Adipocyte
SmoothMuscle
Leukemia_promyelocytic-HL-60
Pathways
BioCarta 2016
CDK Regulation of DNA Replication_Homo sapiens_h_mcmPathway
Oxidative Stress Induced Gene Expression Via Nrf2_Homo sapiens_h_arenrf2Pathway
Agrin in Postsynaptic Differentiation_Homo sapiens_h_agrPathway
Cycling of Ran in nucleocytoplasmic transport_Homo sapiens_h_ranPathway
Prion Pathway_Homo sapiens_h_prionPathway
NCI-Nature 2016
Validated targets of C-MYC transcriptional activation_Homo sapiens_61d3b115-6196-11e5-8ac5-06603eb7f303
Beta3 integrin cell surface interactions_Homo sapiens_c2800165-618d-11e5-8ac5-06603eb7f303
Beta1 integrin cell surface interactions_Homo sapiens_2fd0bc63-618d-11e5-8ac5-06603eb7f303
Signaling events mediated by Hepatocyte Growth Factor Receptor (c-Met)_Homo sapiens_ac39d2b9-6195-11e5-8
BARD1 signaling events_Homo sapiens_75b04491-618c-11e5-8ac5-06603eb7f303
PPI Hub Proteins
CDK2
MAPK14
MYC
MCC
PCNA
Drug Response
RNA-Seq Disease Gene and Drug Signatures from GEO
Erlotinib_11-18_GSE65420_down
PBS-1086_&_Erlotinib_11-18_GSE65420_down
PBS-1086_11-18_GSE65420_down
Panobinostat_JJN3_24hr-10nM_GSE56623_down
Panobinostat_+_5-Azacytidine_JJN3_24hr-10nM_GSE56623_down
Drug Perturbations from GEO down
cetuximab DB00002 human GSE61495 sample 3133
decitabine DB01262 human GSE29077 sample 2539
Curcumin 969516 human GSE16160 sample 3425
decitabine DB01262 human GSE29077 sample 2540
imatinib DB00619 human GSE24493 sample 2764
Transcription
ChEA 2016
E2F7_22180533_ChIP-Seq_HELA_Human
EKLF_21900194_ChIP-Seq_ERYTHROCYTE_Mouse
MYC_18555785_ChIP-Seq_MESCs_Mouse
AR_21909140_ChIP-Seq_LNCAP_Human
NUCKS1_24931609_ChIP-Seq_HEPATOCYTES_Mouse
ENCODE and ChEA Consensus TFs from ChIP-X
MAX_ENCODE
E2F6_ENCODE
MYC_ENCODE
USF1_ENCODE
NFE2L2_CHEA
ENCODE TF ChIP-seq 2015
E2F1_HeLa-S3_hg19
MAX_HCT116_hg19
MYC_A549_hg19
MAX_endothelial cell of umbilical vein_hg19
MYC_HeLa-S3_hg19

Figure 10?

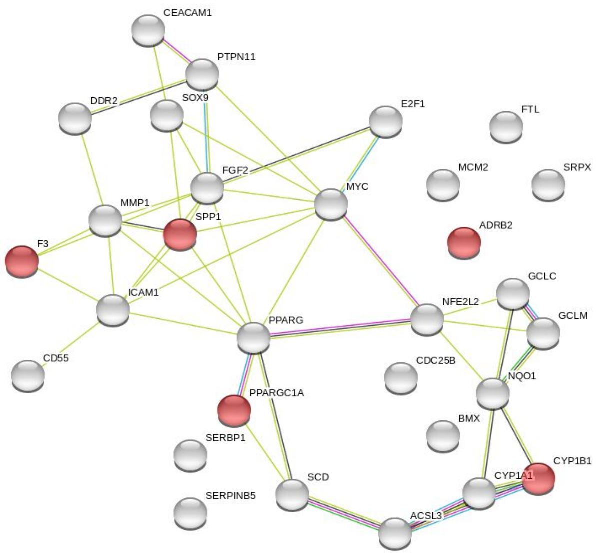

CEACAM1
PTPN11
DDR2
SOX9
E2F1
FTL
FGF2
MCM2
SRPX
MYC
MMP1
SPP1
ADRB2
F3
GCLC
ICAM1
NFE2L2
GCLM
PPARG
CD55
CDC25B
NQO1
PPARGC1A
BMX
SERBP1
CYP1B1
CYP1A1
SCD
SERPINB5
ACSL3


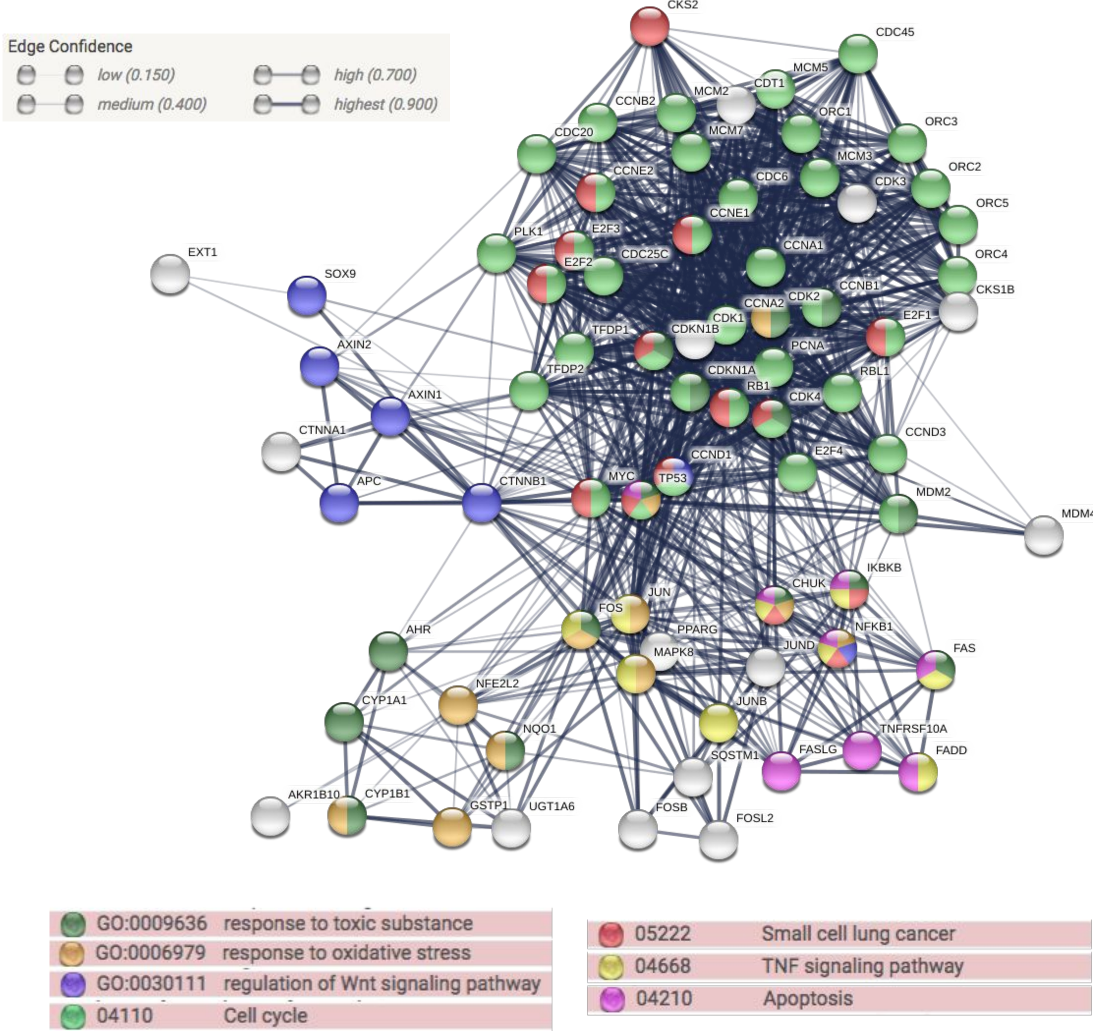

Edge Confidence
low (0.150)
medium (0.400)
high (0.700)
highest (0.900)
CKS2
CDC45
MCM5
CDT1
MCM2
CCNB2
ORC1
ORC3
MCM7
CDC20
MCM3
ORC2
CCNE2
CDC6
CDK3
ORC5
CCNE1
EXT1
PLK1
E2F3
ORC4
CCNA1
SOX9
E2F2
CDC25C
CKS1B
CCNB1
CCNA2
CDK2
CDK1
E2F1
TFDP1
CDKN1B
AXIN2
PCNA
TFDP2
CDKN1A
RBL1
RB1
AXIN1
CDK4
CTNNA1
CCND3
E2F4
CCND1
APC
CTNNB1
MYC
TP53
MDM2
IKBKB
CHUK
JUN
FOS
AHR
PPARG
NFKB1
MAPK8
JUND
FAS
NFE2L2
CYP1A1
JUNB
NQO1
TNFRSF10A
SQSTM1
FASLG
FADD
AKR1B10
CYP1B1
GSTP1
UGT1A6
FOSB
FOSL2
GO:0009636 response to toxic substance
GO:0006979 response to oxidative stress
GO:0030111 regulation of Wnt signaling pathway
04110 Cell cycle
05222 Small cell lung cancer
04668 TNF signaling pathway
04210 Apoptosis
